# Nuclear release of eIF1 globally increases stringency of start-codon selection to preserve mitotic arrest physiology

**DOI:** 10.1101/2024.04.06.588385

**Authors:** Jimmy Ly, Kehui Xiang, Kuan-Chung Su, Gunter B. Sissoko, David P. Bartel, Iain M. Cheeseman

## Abstract

Regulated start-codon selection has the potential to reshape the proteome through the differential production of uORFs, canonical proteins, and alternative translational isoforms. However, conditions under which start-codon selection is altered remain poorly defined. Here, using transcriptome-wide translation initiation site profiling, we reveal a global increase in the stringency of start-codon selection during mammalian mitosis. Low-efficiency initiation sites are preferentially repressed in mitosis, resulting in pervasive changes in the translation of thousands of start sites and their corresponding protein products. This increased stringency of start-codon selection during mitosis results from increased interactions between the key regulator of start-codon selection, eIF1, and the 40S ribosome. We find that increased eIF1-40S ribosome interactions during mitosis are mediated by the release of a nuclear pool of eIF1 upon nuclear envelope breakdown. Selectively depleting the nuclear pool of eIF1 eliminates the changes to translational stringency during mitosis, resulting in altered mitotic proteome composition. In addition, preventing mitotic translational rewiring results in substantially increased cell death and decreased mitotic slippage following treatment with anti-mitotic chemotherapeutics. Thus, cells globally control translation initiation stringency with critical roles during the mammalian cell cycle to preserve mitotic cell physiology.

---

Human cells create extensive proteomic diversity through the differential decoding of the genetic material by alternative transcription, splicing, and translation ^1^. Translational regulation allows cells to rapidly change the set of proteins that are being generated without waiting for changes to the transcriptome, which is particularly important under conditions where transcription is attenuated ^2,3^. Since new transcription is globally inhibited during mitosis ^4-6^, we first assessed the requirement of new translation for mitotic cell viability. Under optimal conditions in cultured human cells, mitosis lasts for ∼60 minutes. However, a key feature of eukaryotic mitosis in both unicellular and multicellular organisms is the ability of cells to sense mitotic errors and delay mitotic progression to ensure sufficient time to correct these errors before undergoing chromosome segregation ^7-10^. We hypothesized that translation would be particularly important under conditions in which cells spend extended periods in mitosis. To test this, we treated cells with the KIF11 inhibitor, S-trityl-L-cysteine (STLC) ^11^ to induce a mitotic delay through the conserved spindle assembly checkpoint (Fig. 1A). Strikingly, we found that mitotically-arrested cells were exquisitely sensitive to cycloheximide (CHX, a translational inhibitor), undergoing massive cell death after only 4 hours of cycloheximide treatment (Fig. 1B and 1C). This is in stark contrast to interphase cells, which can survive in presence of translational inhibition for up to 2 days in culture (Extended data Fig. 1A and 1B) ^12^. The cycloheximide-induced death for mitotically-arrested cells was partially suppressed by the addition of the proteasome inhibitor MG132 to inhibit protein degradation (Fig. 1B and 1C) suggesting that protein turnover increases the requirement for new protein synthesis. These results suggest that there is a strong requirement for new translation when cells experience a mitotic delay.

**Figure 1.**
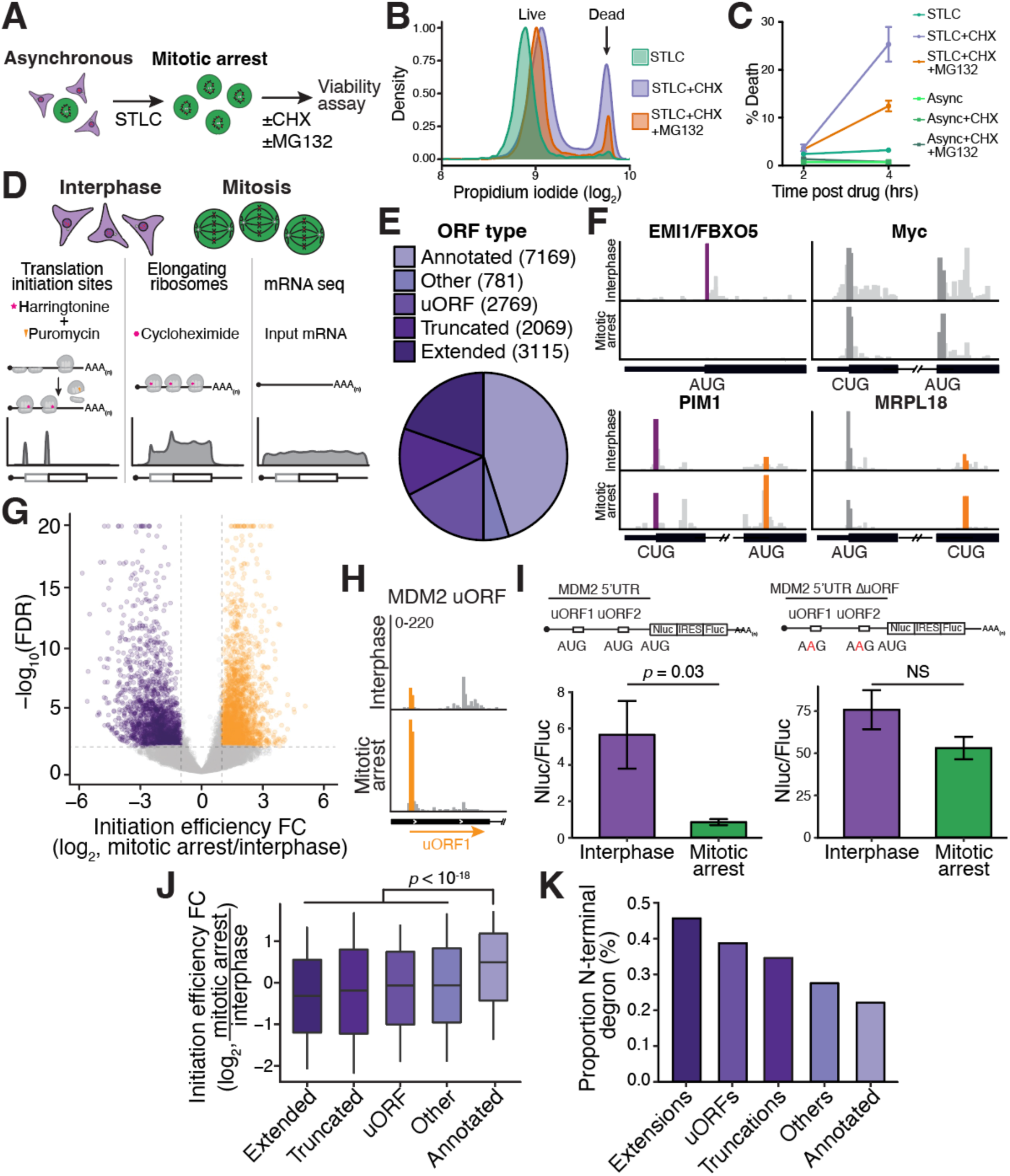
Thousands of translation initiation sites change in efficiency during mitosis. (**A**) Schematic of mitotic cell viability assay using propidium iodide staining in the presence of cycloheximide (CHX) ± MG132. (**B**) Representative distribution of live or dead mitotically arrested cells treated with CHX or CHX+MG132 for 4-hours. (**C**) Quantification of cell viability 2- and 4-hour post CHX or CHX+MG132 addition. N = 3 biological replicates. (**D**) Schematic outline of translation initiation site sequencing, ribosome profiling, and paired input RNA sequencing showing hypothetical read distributions for an mRNA with two translation initiation sites. (**E**) Distribution of ORF types identified using RiboTISH. (**F**) Read distribution from translation initiation site sequencing surrounding the start codon for the indicated established alternative translation isoforms from interphase and mitotic cells. Purple indicates initiation site that is preferentially repressed while orange is preferentially translated during mitosis. Dark grey - initiation sites that don’t change significantly. Light grey - non-translation initiation site reads. Significance cutoff described in (G). (**G**) Volcano plot of translation site differences between interphase and mitotically-arrested cells. Each point represents a translation initiation site. Orange – significantly upregulated sites in mitosis. Purple - significantly downregulated sites. Translation initiation sites with FDR < 0.001 and a fold change > 2 were called as significant. (**H**) Read distribution from translation initiation site sequencing surrounding the MDM2 uORF1, which is preferentially translated during mitosis. (**I**) Top, schematic outline of luciferase mRNA reporters to test MDM2 translation in the presence (left) or absence (right) of the MDM2 uORFs. Bottom, normalized nano/firefly luciferase activity from interphase or mitotically-arrested cells, 3 hours post mRNA transfection. N = 3 biological replicates. Statistics reflect an unpaired student’s T-test. (**J**) Boxplot of translation initiation efficiency fold change between mitotic arrest and interphase (Y-axis) and the class of ORF (X-axis). Statistics indicate a Mann-Whitney U-test. (**K**) Proportion of predicted N-terminal degrons (where the second amino acid is glycine, lysine, arginine, or cysteine) in different ORF classes.

## Thousands of translation initiation sites change in efficiency during mitosis

As translation is important for maintaining cell viability during a mitotic delay, we next sought to assess the translational landscape during mitosis. Differences in start-codon selection though leaky ribosome scanning, internal ribosome entry, and initiation at near-cognate start codons can each result in the usage of multiple translation initiation sites to generate multiple protein products from a single mRNA ^1,13^ ^14^. To assess cell cycle-dependent changes in start-codon selection, we performed translation initiation-site sequencing, a modified ribosome profiling-based strategy to enrich for initiating 80S ribosomes (Fig. 1D; see Methods). By performing translation initiation-site sequencing, ribosome profiling, and RNA-seq in parallel, we were able to identify transcriptome-wide changes in translational start site usage between interphase and mitosis (Extended data Fig. 1C-G) ^15^. To quantitatively interrogate the relative efficiency of each translation initiation site, we computed the translation initiation efficiency corresponding to the read counts at an initiation site normalized to mRNA abundance (Extended data Fig. 1C; see Methods). In particular, we assessed the translation initiation profile of interphase cells, cycling mitotic cells, and cells arrested in mitosis by treatment with the kinesin-5 inhibitor, STLC (Extended data Fig. 1H). Nocodazole treatment has been shown to activate the integrated stress response ^16^. In contrast, these mitotic synchronization strategies did not activate stress responses, as monitored by the lack of eIF2α phosphorylation and the absence of changes to ATF4 translation (Extended data Fig. 1A and 1I). This allowed us to test cell cycle-specific differences in the absence of confounding factors. Translation initiation site profiles were highly correlated between cycling mitotic cells and cells arrested in mitosis using STLC (Extended data Fig. 1J and 1K), indicating that the observed changes in translation initiation efficiency represent the mitotic state rather than reflecting a specific synchronization strategy.

From our combined datasets, we identified a total of 15902 translation initiation sites from 7734 different genes (Fig. 1E; Table S1). These initiation sites fell into five open reading frame (ORF) categories: previously annotated ORFs (7169), in-frame N-terminal extensions (3115), in-frame N-terminal truncations (2069), upstream ORFs (uORFs; 2769), and other ORFs (either internal-out-of-frame ORFs or downstream ORFs present in the 3’-UTR; 781) (Fig. 1E). These identified start sites include many previously established non-canonical translation initiation sites. For example, we detected alternative translation initiation sites for BAG1 ^17^, POLG ^18^, and the oncogene c-Myc ^19^, amongst others (Fig. 1F; Table S1).

To assess changes to translation start site usage, we compared the interphase and mitotic translation initiation efficiencies for each start site (Methods; ^15^). We identified 3993 translation initiation sites that were differentially used between interphase and mitotically arrested cells (> 2-fold change and FDR < 0.01) (Fig. 1G; Table S1). Importantly, our data accurately captured the expected cell-cycle dependent translational repression of EMI1/FBXO5 ^20^ (Fig. 1F) and the mitotic activation of translation for TOP motif-containing mRNAs ^21^ (Extended data Fig. 1A; Table S1). The changes in translation initiation between interphase and mitosis also alter the expression of multiple N-terminal protein isoforms. For example, we observed substantial changes in the relative ratios of the previously identified alternative translational isoforms for PIM1 ^22^ and MRPL18 ^23^, amongst others (Fig. 1F; Table S1).

In addition to alternative translational isoforms, we observed striking changes in uORF translation. For example, we found that the uORFs in the MDM2 5’UTR ^24^ are preferentially translated during mitosis, whereas translation of the annotated MDM2 protein during mitosis was preferentially repressed ^25^ (Fig. 1H; Table S1, Table S2). As the regulation of MDM2 levels plays a critical role in cell viability following a mitotic arrest ^25^, we sought to test whether preferential MDM2 uORF translation during mitosis acts to reduce MDM2 protein synthesis. To test this, we generated luciferase mRNA reporters harboring different versions of the MDM2 5’ UTR (Fig. 1I). The wild-type MDM2 5’ UTR reporter was repressed 6.7-fold in mitosis relative to interphase. In contrast, a reporter in which the uORF start-codons are eliminated was only repressed 1.4x during mitosis, suggesting that differential cell cycle uORF translation controls MDM2 protein synthesis (Fig. 1I), highlighting the roles in altered start-codon selection for mitotic translational control.

In addition to changes in specific initiation sites, we observed global changes in ORF type usage during mitosis in which previously unannotated alternative translational isoforms (uORFs, N-terminal truncations, extensions, and internal ORFs) were preferentially repressed in mitotic cells relative to annotated ORFs (Fig. 1J). Such altered translational regulation would be particularly impactful for its ability to remodel steady-state protein levels for short-lived proteins (<8 hours), which constitute ∼10% of the proteome ^26^. For example, prior work found that combined Emi1 translational repression and protein destabilization during mitosis are required to control Emi1 levels for accurate cell division ^20^. Therefore, we considered whether proteins that are preferentially repressed during mitosis are enriched for short-lived proteins. The identity of a protein’s N-terminal amino acid affects its protein stability by acting as an N-terminal degron ^27^. We found that annotated ORFs, which are preferentially translated during mitosis (Fig. 1J), are depleted for N-terminal destabilizing degrons, potentially contributing to protein stability (Fig. 1K). In contrast, we found that alternative N-terminal protein extensions were enriched for destabilizing degrons (Fig. 1K) together with being preferentially repressed during mitosis (Fig. 1J). We propose that the combined action of translational control and post-translational protein stability of annotated and alternative ORFs contributes to reshaping the mitotic proteome. Together, our results demonstrate that there is a widespread change in the translational landscape during mitosis, with the altered usage of thousands of translation initiation sites.

## Non-AUG translation is preferentially repressed during mitosis

To understand the mechanistic basis for the widespread changes in translation initiation, we generated a linear regression model to identify sequence features that contribute to the change in translation between mitosis and interphase (see Methods; Extended data Fig. 2A). We found that start-codon identity was one of the most important features for determining changes in translation initiation efficiency between interphase and mitosis (Fig. 2A). Our results suggest that AUG initiation sites are preferentially used during mitosis compared to non-AUG initiation sites (Fig. 2B and 2C). This preferential repression of non-AUG initiation sites likely explains the mitotic repression of previously unannotated ORFs, ∼80% of which initiate from near-cognate start codons (Extended data Fig. 2B).

**Figure 2.**
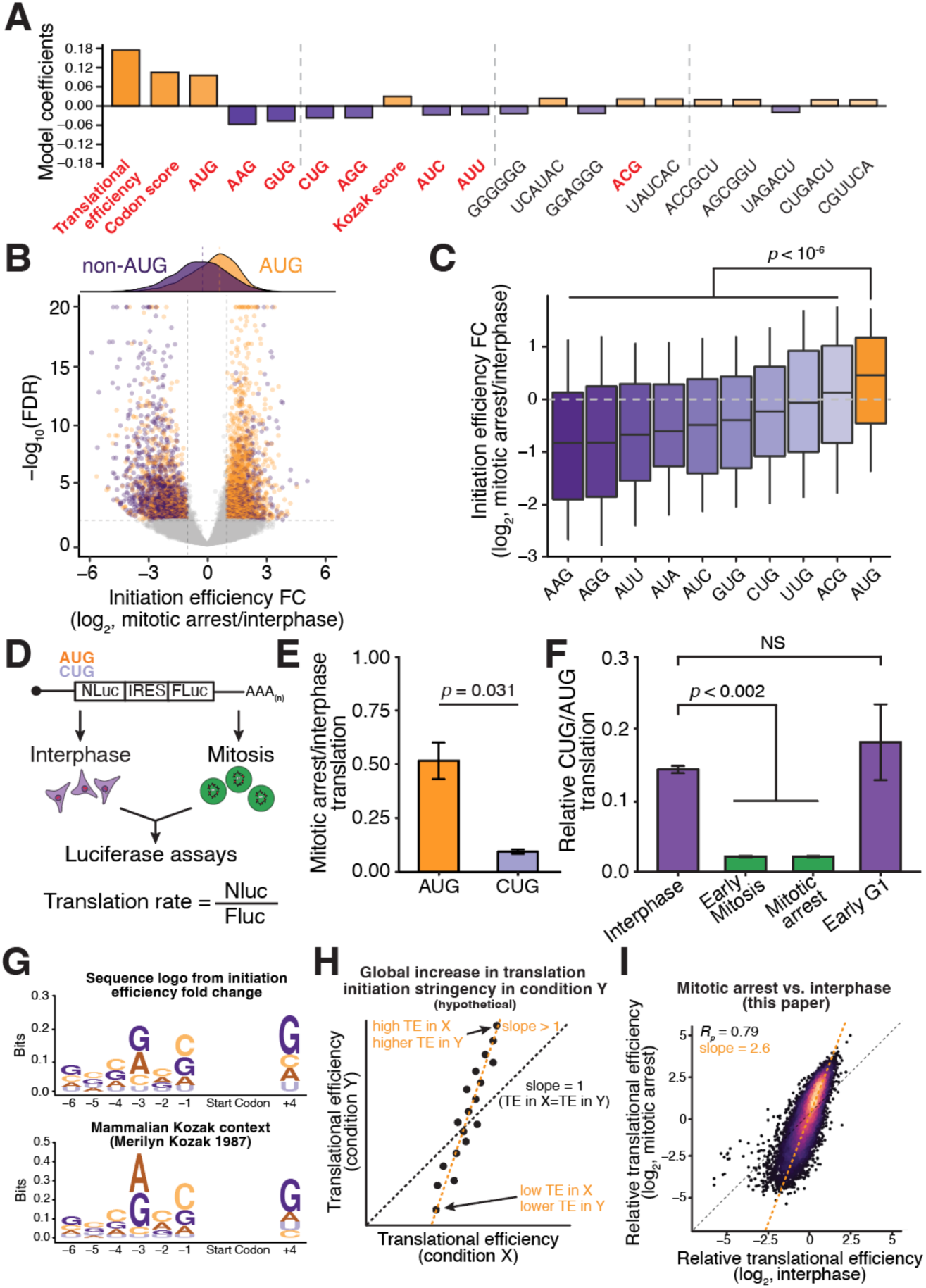
Translation initiation stringency increases during mitosis. (**A**) Graph showing coefficients from a linear regression model to predict initiation efficiency fold change between mitotic arrest and interphase. Coefficients were averaged among 10 models. The top 20 features are shown, with features related to the start codon highlighted in red. (**B**) Volcano plot of translation site differences between mitotic and interphase cells. Each point represents a translation initiation site, with orange points representing significantly different AUG initiation sites and purple representing significantly different non-AUG initiation sites. Translation initiation sites with FDR <0.001 and >2 fold change were called as significant. (**C**) Boxplot of translation initiation efficiency fold changes between mitotic arrest and interphase (Y-axis) and the start codon (X-axis). (**D**) Schematic of mRNA luciferase reporter used to compare AUG and CUG translation in different cell cycle stages. Normalizing Nano luciferase to Firefly luciferase activity accounts for mRNA abundance and transfection efficiency. (**E**) Graph showing translation rates for AUG and CUG mRNA reporters during an 8-hour mitotic arrest normalized to interphase cells. Error bar represents standard error of the mean, N = 3 biological replicates. Statistics reflect an unpaired student’s T-test. (**F**) Graph showing differences in the CUG and AUG luciferase reporters (from D) in interphase cells, cells arrested in mitosis for 1 hour (early mitosis), 8 hours (mitotic arrest), or cycling G1 cells. Error bar represents standard error of the mean, N = 2 biological replicates and unpaired student’s T-test used for this data. (**G**) Top, sequence logo created using weighted translation initiation efficiency fold changes between mitotic arrest and interphase. Sequence logo is similar to the known strong Kozak context. Bottom, unweighted sequence logo of nucleotides flanking the start codon for annotated ORFs as described in ^29^. (**H**) Schematic representation showing global increase in translation initiation stringency in condition on the Y-axis relative to X-axis. A slope >1 is suggestive of an increase translation initiation stringency. (**I**) Relative translational efficiency correlations between interphase and mitotically-arrested cells indicates increased translation initiation stringency during mitosis.

As an orthogonal method to assess the preferential repression of non-AUG translation initiation sites during mitosis, we generated luciferase mRNA-translation reporters containing either an AUG or CUG start-codon and used a normalized luciferase signal as a proxy for translation initiation efficiency (Fig. 2D). The translational efficiency of the AUG reporter was reduced 2-fold during mitosis, consistent with prior work ^20^. However, the translational efficiency of the CUG reporter was reduced 10-fold during mitosis, 5-fold greater repression than the AUG reporter, consistent with our ribosome profiling analysis (Fig. 2E). This preferential repression of the CUG luciferase reporter was observed during both a short (1 hour) and prolonged (8 hour) mitotic arrest (Fig. 2F; Extended data Fig. 2C). In addition, the CUG luciferase reporter displayed preferential repression during mitosis independent of the synchronization strategy used to isolate mitotic cells, as we detected a similar repression in both STLC and taxol-arrested cells (Extended data Fig. 2C). Moreover, the CUG reporter was not repressed in cells present in late G2 or early G1 (Fig. 2F; Extended data Fig. 2C), indicating that the repression of non-AUG initiation sites was restricted to mitosis. Non-AUG translation was also preferentially repressed during mitosis in non-transformed human RPE1 cells and mouse macrophage cells (RAW264.7) (Extended data Fig. 2C-E) indicating that increased translational stringency during mitosis is conserved between human and mouse cells and is not restricted to a specific cell type or transformation state.

Together, our results reveal a conserved mitotic translational program in which translation initiation is strongly biased towards AUG translation initiation sites. In contrast, interphase translation is more permissive, allowing increased usage of near-cognate initiation sites.

## Translation initiation stringency is increased during mitosis

In addition to start-codon identity, the Kozak context had an important effect on cell cycle start-codon usage in our linear regression model (Fig. 2A). Thus, we next sought to assess whether weaker translation initiation sites, independent of the triplet start-codon, are preferentially repressed in mitosis. Indeed, we observed that the optimal vertebrate Kozak sequence ^28,29^ was associated with preferential translation during mitosis whereas ORFs with a poor Kozak-context were preferentially repressed (Fig. 2G; Extended data Fig. 3A). Together, our analysis indicated that the relative level of translation initiation efficiency observed for a start-codon during interphase can explain 67% of the repression of non-AUG initiation sites during mitosis (Extended data Fig. 3B). This preferential repression of weak initiation sites coupled with the preferential usage of strong sites is indicative of increased translation initiation stringency during mitosis.

**Figure 3.**
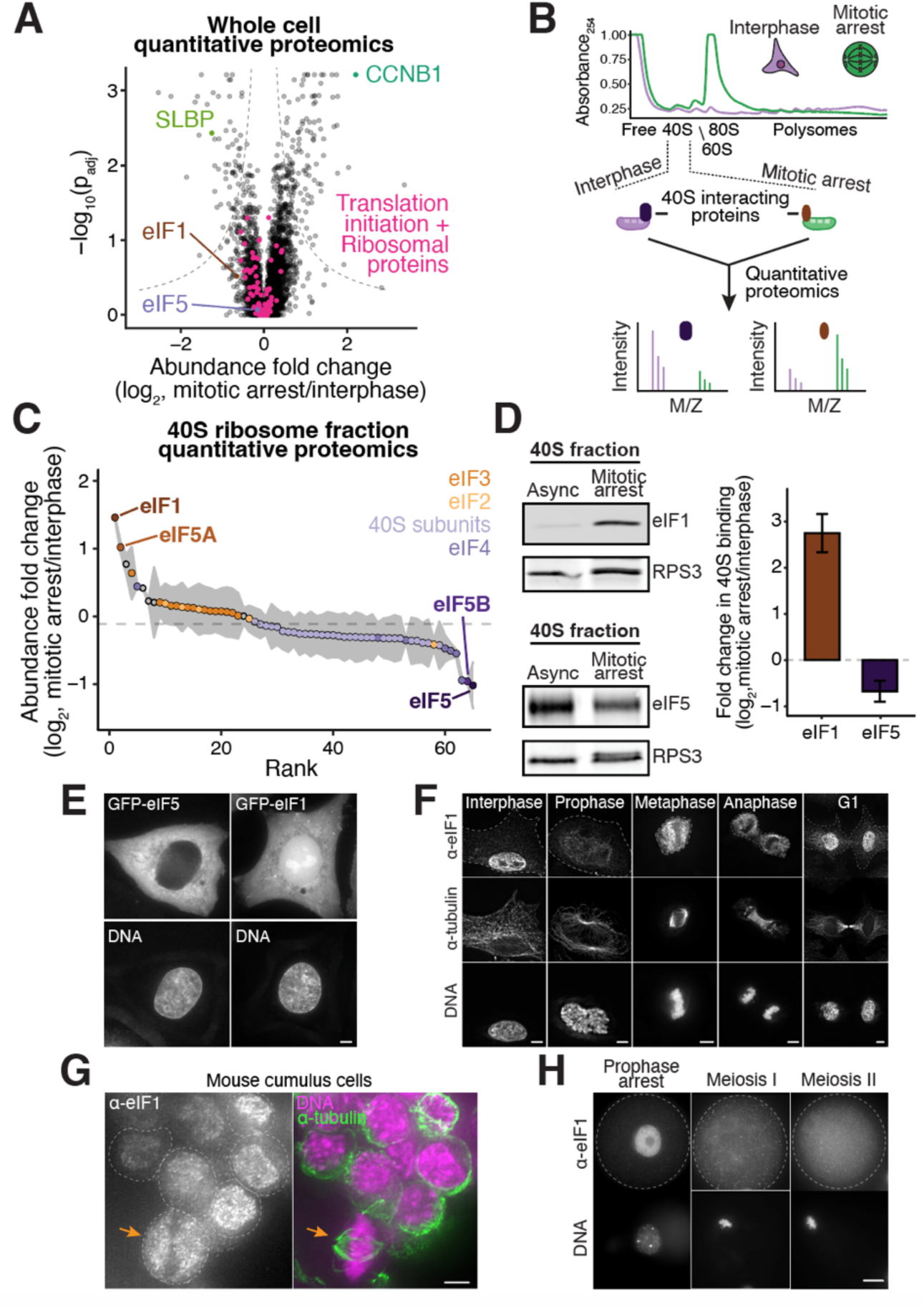
Nuclear release of eIF1 increases ribosome association during mitosis. (**A**) Volcano plot showing whole cell quantitative proteomics from interphase and mitotically arrested HeLa cells. CCNB1 and SLBP are positive controls, as they are known to be increased and decreased, respectively, during mitosis. The steady state protein levels of translation initiation factors and ribosomal proteins don’t significantly change between interphase and mitotic arrest. Dashed lines indicate the significance threshold. N = 3 biological replicates. (**B**) Schematic of ribosome fractionation and quantitative mass spectrometry analysis to identify proteins with differential ribosome associations between interphase and mitotic arrest. Cell extracts from interphase and mitotic cells were separated on a sucrose gradient. Free, 40S, 60S, and 80S ribosome fractions were isolated for quantitative mass spec analysis. (**C**) Plot showing rank-ordered fold change from quantitative proteomic analysis of the 40S ribosome fraction from interphase and mitotic cells. The association of most translation initiation factors and ribosomal proteins does not change in the 40S fraction during mitotic arrest, whereas eIF1 preferentially associates and eIF5 shows decreased association with the 40S ribosome. Each point represents the average of 2 biological replicates and the shading indicates standard error of mean. (**D**) Western blot analysis of 40S ribosome fractions isolated from interphase and mitotic cells (left). Quantification of western blots showing the change in eIF1/RPS3 or eIF5/RPS3 ratio between interphase and mitotic arrest (right, error bar represents standard error of the mean, N = 2 biological replicates). (E) Live-cell imaging of GFP-eIF5 (left) and GFP-eIF1 (right). GFP-eIF5 displays largely cytoplasmic localization whereas GFP-eIF1 shows cytoplasmic, nuclear, and nucleolar localization. Scale bar, 5 µm. (F) Immunofluorescence of endogenous eIF1 (anti-eIF1) in HeLa cells throughout the cell cycle. α-eIF1 image intensity is scaled identically between images. α-tubulin and DNA image intensity were adjusted in each cell to highlight cell morphology. Dotted lines represent cell boundaries. Scale bar, 5 µm. (**G**) Immunofluorescence showing endogenous eIF1 localization (anti-eIF1) in interphase and mitotic mouse cumulus cells. Yellow arrow indicates mitotic cumulus cell. Dotted lines represent cell boundaries. Scale bar, 5 µm. (**H**) Immunofluorescence showing endogenous eIF1 localization in mouse GV, meiosis I, and meiosis II eggs. Dotted lines represent cell boundaries. Scale bar, 20 µm.

As an orthologous strategy to assess changes in translational stringency during mitosis, we analyzed translational efficiency (which measures elongating ribosomes instead of only initiating ribosomes), a well-established metric to measure translation output ^30,31^ (Fig. 2H). When comparing the translational efficiency of annotated ORFs between interphase and mitotic arrest, we observed the preferential mitotic repression of initiation sites with low interphase translational efficiency, consistent with an increase in translational stringency during mitosis (Fig. 2I; Extended data Fig. 3C). Reanalysis of previously published cell cycle-resolved ribosome profiling data ^21^ revealed a similar increase in translational stringency during mitosis, relative to asynchronous or S-phase arrested cells (Extended data Fig. 3D-F). This enhanced mitotic translation initiation stringency cannot be explained due to the global attenuation of translation during mitosis, as analysis of conditions known to inhibit global translation, including hypoxic conditions ^32^, treatment with Torin-1 ^33^, or arsenite ^34^, did not reveal global changes to translation initiation stringency (Extended data Fig. 3G-I). Based on the effects of hypoxia, Torin-1, and arsenite treatments, these data also suggest that global translational repression due to eIF2α phosphorylation or 4E-BP dephosphorylation is not sufficient to induce an increase in translation initiation stringency.

Collectively, our results reveal a global rewiring of translation initiation during mitosis. Mitotic translation initiation is strongly biased towards strong translation initiation sites, such as AUG start codons and optimal Kozak context start sites. This enhanced translational stringency alters the relative synthesis of thousands of ORFs, including in-frame N-terminal protein isoforms and uORFs.

## eIF1 preferentially associates with the ribosome during mitosis

As our results indicated that translation initiation stringency changes markedly throughout the cell cycle, we hypothesized that there may be differences in the translation initiation machinery between interphase and mitosis ^35^. Using quantitative whole cell proteomics, we found that the levels of translation factors and ribosome proteins were similar between interphase and mitotic arrest (Fig. 3A; Table S3), suggesting that changes in translation initiation factor expression is not likely to explain differences in translational control. Thus, we next evaluated the changes in ribosome association between interphase and mitosis arrest using sucrose gradient fractionation and quantitative mass spectrometry (Fig. 3B; Extended data Fig. 4A-B, Table S4). Most translation initiation factors did not display clear changes in their association with the 40S or 60S ribosome fractions when comparing interphase and mitotic cells (Fig. 3C; Extended data Fig. 4C). In contrast, eIF1 and eIF5A were substantially enriched in the 40S fraction during mitosis relative to interphase, whereas eIF5 and eIF5B were depleted (Fig. 3C).

**Figure 4.**
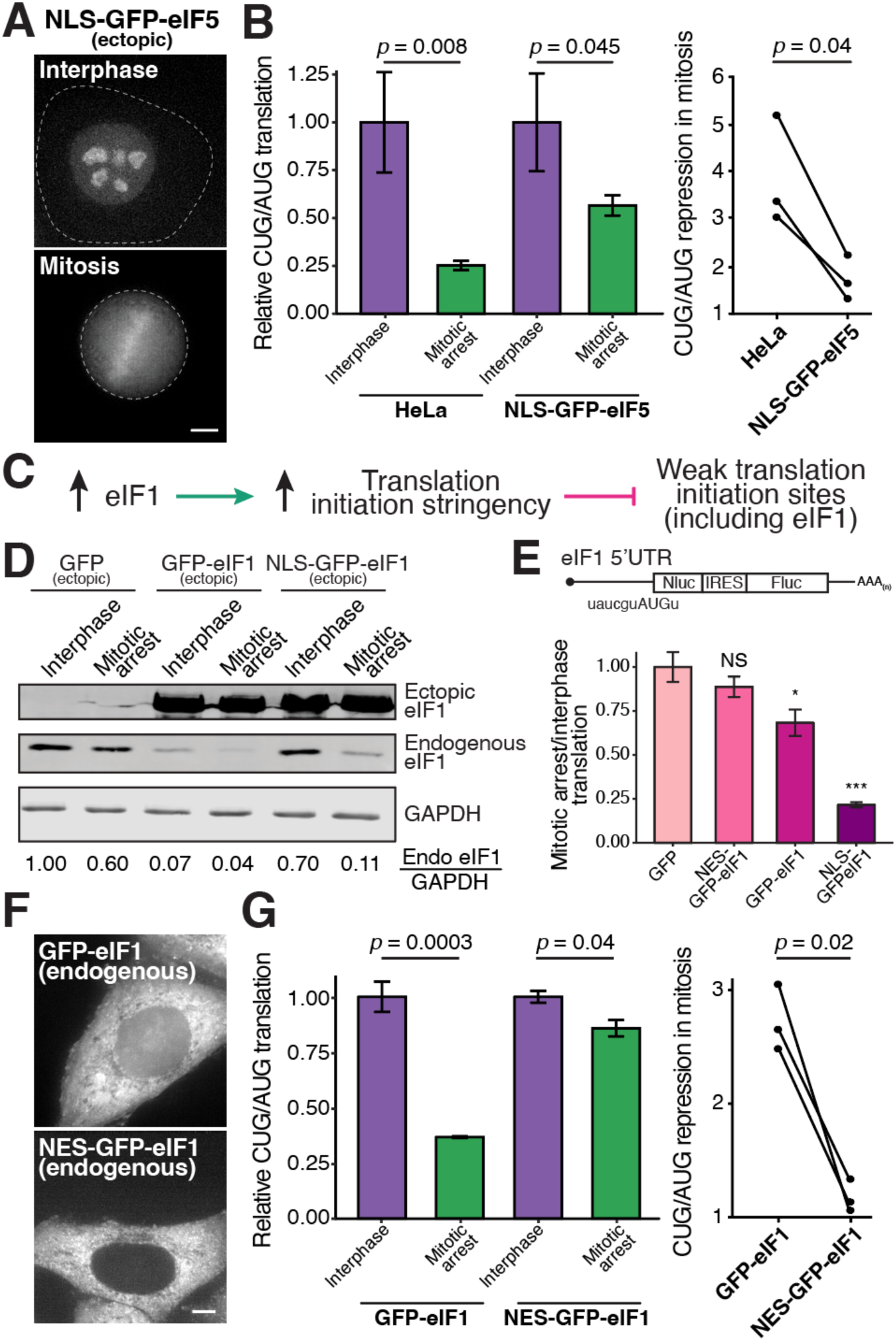
Nuclear/cytoplasmic ratios of initiation factors differentially control translation initiation stringency during mitosis. (**A**) Live cell imaging showing localization of NLS-eIF5-GFP-NLS during interphase and mitosis. Dotted lines represent cell boundaries. Scale bar, 5 µm. (**B**) Graph showing quantification of luciferase reporters in the indicated conditions. The CUG luciferase reporter is repressed 3.9-fold over the AUG reporter in HeLa cells during mitosis and 1.7-fold in cells expressing nuclear-specific eIF5. Mitotic CUG/AUG translation was normalized to interphase within each cell line. N = 3 biological replicates. Unpaired student’s T-test was used for plots on the left. Paired T-test was used for plots on the right. (**C**) Proposed model for eIF1 autoregulatory loop as described in ^40^. Increased eIF1 expression suppresses eIF1 translation. (**D**) Western blot showing ectopic and endogenous eIF1 levels. Ectopic expression of GFP-eIF1 induces a ∼10-fold decrease in endogenous eIF1 levels in both interphase and mitotic cells. Ectopic expression of NLS-GFP-eIF1 specifically induces a ∼10-fold decrease in endogenous eIF1 protein levels in mitosis but only modestly affects interphase eIF1 levels. (**E**) Graph showing translational changes in luciferase reporter harboring eIF1 5’UTR. Luciferase activity from wild-type eIF1 5’UTR reporter (weak Kozak, uaucguAUGu) was normalized an eIF1 5’UTR reporter harboring a strong Kozak context (gccaccAUGg). Ectopic expression of NLS-GFP-eIF1 preferentially represses the translation of a reporter harboring the eIF1 5’UTR (weak Kozak context) during mitosis. N = 3 biological replicates. Statistics reflect an unpaired student’s T-test compared to ectopic GFP expression, * p < 0.05, *** p < 0.0005. (**F**) Live cell imaging showing localization of endogenous GFP-eIF1 and NES-GFP-eIF1 during interphase. Scale bar, 5 µm. (**G**) Graph showing quantification of luciferase reporters in the indicated conditions. The CUG reporter is repressed 2.7-fold over AUG reporter in control GFP-eIF1 cells during mitosis and 1.2-fold in cytoplasmic-specific eIF1 cells. Mitotic CUG/AUG translation was normalized to interphase within each cell line. Error bar represents standard error of the mean, N = 3 biological replicates and Unpaired student’s T-test was used for plots on the left. Paired T-test was used for plots on the right.

eIF1, eIF5A, eIF5, and eIF5B are regulators of start codon selection ^36-44^, with eIF1 acting directly to increase the stringency of start codon selection and eIF5 acting to decrease stringency ^40,42,45^. We chose to focus our analysis on eIF1 and eIF5 based on these established roles in controlling start codon selection and on the magnitude of the changes in their 40S interaction between interphase and mitosis. eIF1 and eIF5 also compete with each other for binding to the 40S ribosome ^40,42,46^, consistent with the opposing changes observed in eIF1-40S and eIF5-40S associations during mitosis (Fig. 3C and 3D). We confirmed the increased ribosome association for eIF1 and decreased association for eIF5 during mitosis by western blotting of 40S fractions (Fig. 3D). Based on this behavior, we propose that altered eIF1 and eIF5 associations with the ribosome contribute to the altered translation initiation stringency during mitosis.

## Nuclear eIF1 is released into the cytoplasm during mitosis and meiosis

We next sought to define the molecular mechanisms responsible for the change in eIF1/eIF5-ribosome association between interphase and mitosis. As we only observed increased translational stringency in mitosis, but not late G2 or early G1 (Fig. 2F; Extended data Fig. 2C), the underlying mechanisms must be rapid and switch-like. As the levels of ribosome components and translation initiation factors protein, including eIF5 and eIF1, are similar throughout the cell cycle (Fig. 3A; Extended data Fig. 4D, and Table S3), a change in protein levels is unlikely to explain the differential ribosome interactions. Similarly, both our analysis and prior work found that the phosphorylation state of eIF1 and eIF5 does not change substantially between interphase and mitosis ^47,48^ (data not shown).

Strikingly, we found that eIF1 and eIF5 display distinct sub-cellular localization, providing a potential explanation for their differential activity in interphase and mitosis. Using N-terminally GFP-tagged fusion proteins, which we found did not disrupt protein activity (Extended data Fig. 5A), we observed that GFP-eIF5 localized exclusively to the cytoplasm (Fig. 3E), similar to most other components of the translation machinery ^49,50^. In contrast, GFP-eIF1 localized to both the cytoplasm and the nucleus during interphase in HeLa cells, untransformed human RPE1 cells, and mouse RAW264.7 cells (Fig. 3E; Extended data Fig. 5B and 5C). Similarly, endogenous eIF1 (detected using an affinity-purified antibody) localized primarily to the nucleus of interphase cells, but was diffuse throughout the cell during mitosis in HeLa cells, mouse 3T3 cells, and ovarian mouse cumulus cells (Fig. 3F, 3G; Extended data Fig. 5D-F). Finally, we tested eIF1 localization in mouse oocytes (Fig. 3H). During mammalian meiosis, oocytes arrest in metaphase II for days or until fertilization occurs with an intact meiotic spindle and a disassembled nuclear envelope. In GV arrested mouse oocytes with an intact nucleus, endogenous eIF1 localized primarily to the nucleus (Fig. 3H). However, upon meiotic maturation, we observed the nuclear release of eIF1 in meiosis I which persists throughout meiosis II. We propose that this substantial nuclear pool of eIF1 is retained during interphase at least in part through protein-protein interactions. Indeed, based on both GFP-eIF1 and endogenous anti-eIF1 immunoprecipitations from interphase cells, we found that eIF1 associated with multiple nuclear proteins (Extended data Fig. 6A-C, Table S5, and Table S6). Together, these data suggest that a population of eIF1 is present in the nucleus of interphase cells but is released into the cytoplasm following nuclear envelope breakdown in dividing cells across different mammalian organisms, cell types, and cell states (Extended data Fig. 5G and 5H).

**Figure 5.**
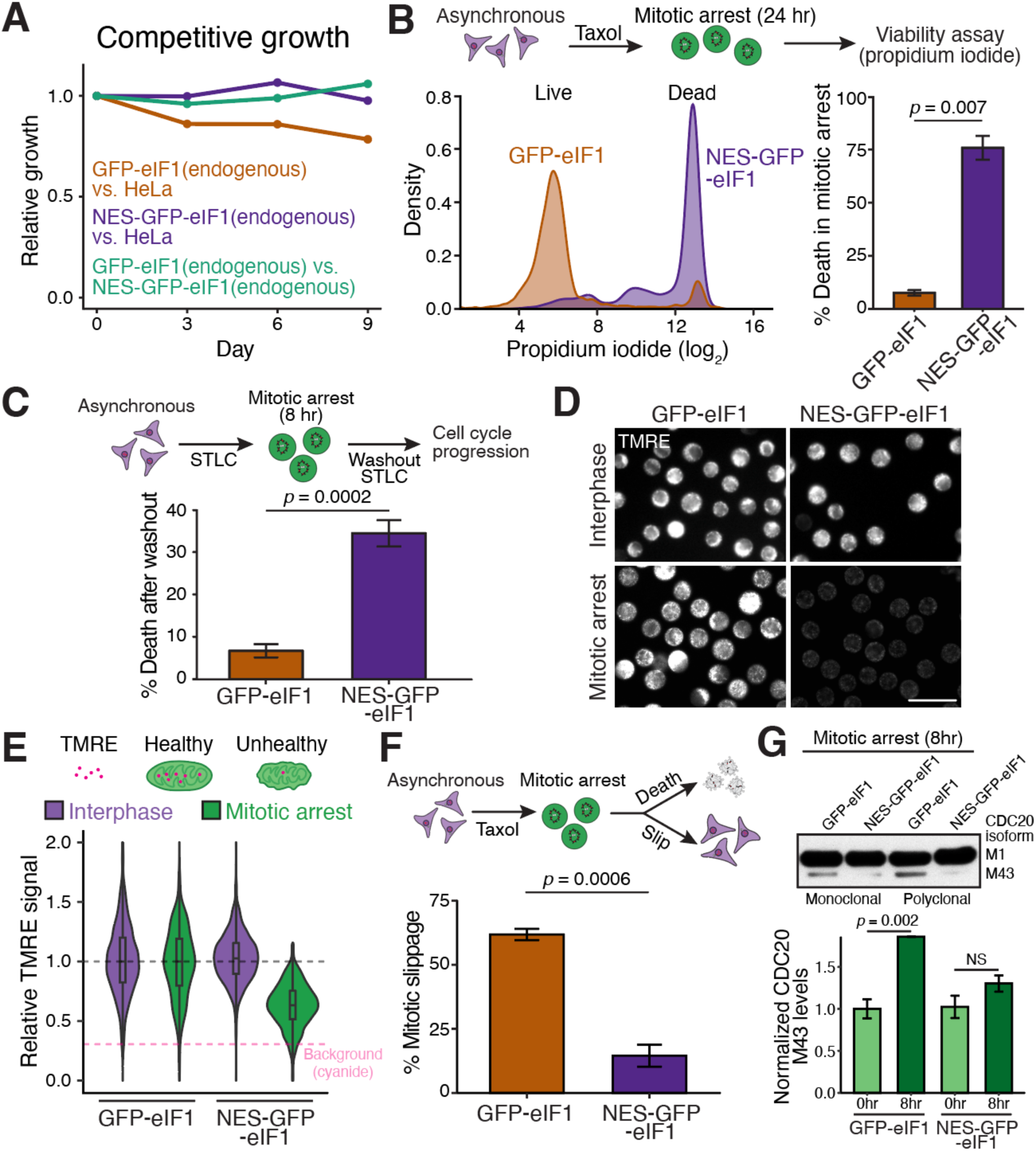
Depleting nuclear eIF1 sensitizes cells to anti-mitotic chemotherapeutics. (**A**) Competitive growth assay showing the relative growth of control HeLa cells and GFP-eIF1 (brown) or NES-GFP-eIF1 (purple), and GFP-eIF1 and NES-GFP-eIF1 (green) over time. (**B**) Top, schematic outline of mitotic death assay. Cells were arrested in mitosis for 24 hours using taxol before measuring viability. Left, representative flow cytometry plot of propidium iodide signal from 24 hour mitotically-arrested GFP-eIF1 or NES-GFP-eIF1. Right, quantification of mitotic death assays. Error bar represents standard error of the mean, N = 3 biological replicates and Unpaired student’s T-test was used for plots on the right. (**C**) Top, schematic outline to test viability after a short mitotic delay. Bottom, graph showing quantification of cell death after STLC washout using live cell imaging. Error bar represents standard error of the mean, N = 3 biological replicates and unpaired student’s T-test was used. Within each biological replicate, >100 cells were quantified. (**D**) Representative image of TMRE staining from interphase and mitotically-arrested endogenous GFP-eIF1 and NES-GFP-eIF1 cells. Scale bar, 50 µm. (**E**) Representative violin plot showing the TMRE signal in interphase (purple) and mitotically arrested (green) endogenous GFP-eIF1 or NES-GFP-eIF1 cells. >1000 cells were quantified per condition. Magenta dotted line represents median TMRE signal upon cyanide (CCCP) treatment. CCCP is an electron transport chain poison, resulting in mitochondrial depolarization therefore decreased TMRE staining and can be used as a marker for no mitochondrial activity. (**F**) Top, schematic showing strategy to test the outcome of mitotically-arrested cells. Bottom, graph showing quantification of the proportion of mitotic slippage during taxol-induced mitotic arrest. Error bar represents standard error of the mean, N = 3 biological replicates and unpaired student’s T-test was used. Within each biological replicate, >100 cells were quantified. (**G**) Top, western blot analysis of CDC20 protein isoform expression in mitotically-arrested endogenous GFP-eIF1 and NES-GFP-eIF1 cells. Bottom, quantification of CDC20 short M43 isoform normalized to GAPDH in early (0 hr) and prolonged (8 hr) mitotic cells from GFP-eIF1 and NES-GFP-eIF1 cells. Nuclear eIF1 is required for expression of CDC20 M43 isoform during mitosis. N = 3 biological replicates and unpaired student’s T-test was used.

**Figure 6.**
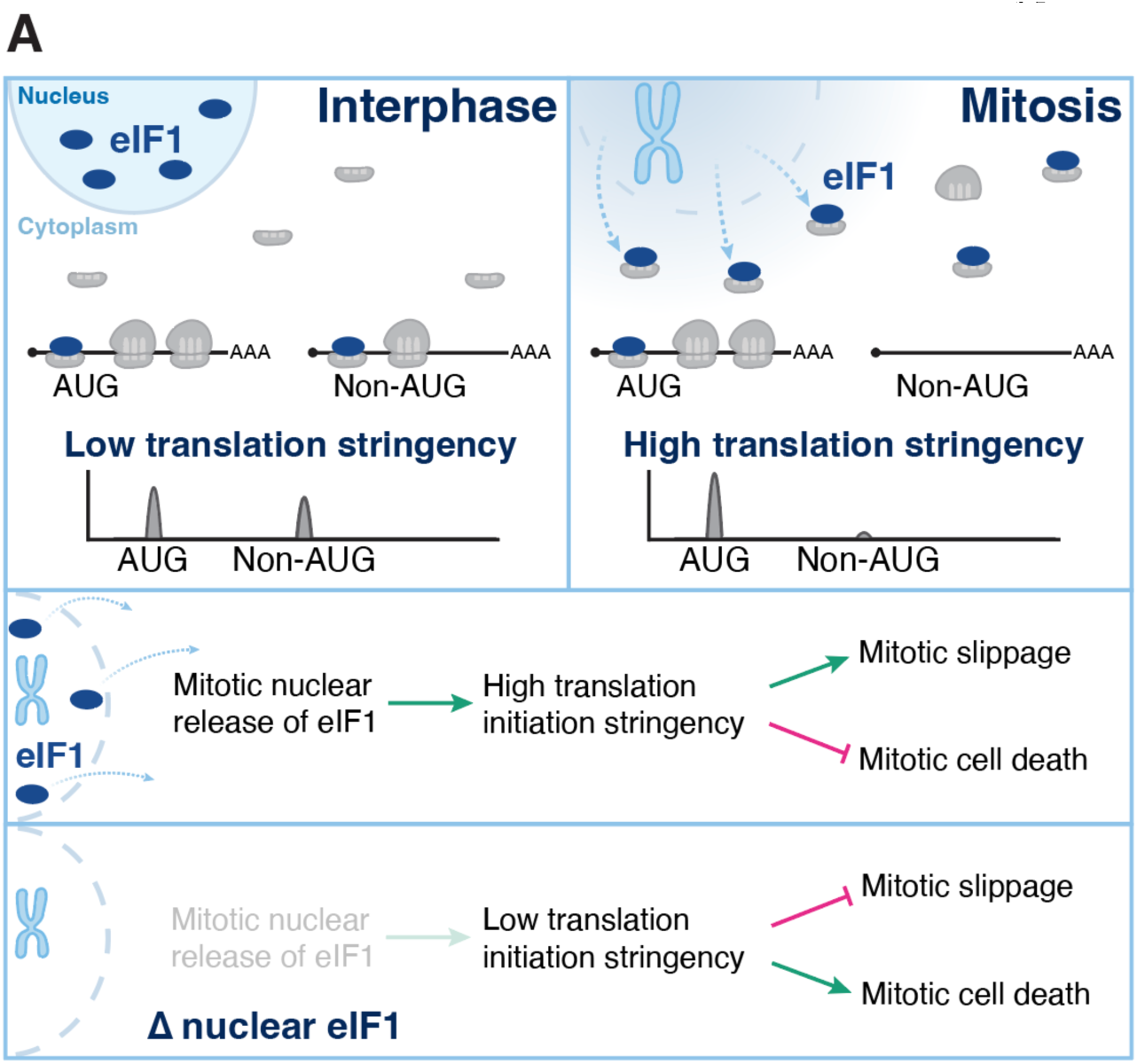
Programmed release of nuclear eIF1 enhances stringency of start codon selection to enable physiological mitotic arrest. (**A**) Schematic of the model of nuclear release of eIF1 and its role in controlling mitotic cell physiology and translation.

## Nuclear/cytoplasmic ratios of initiation factors differentially control translation initiation stringency during mitosis

The increased availability of eIF1 molecules in the cytoplasm following nuclear release during mitosis might act to mimic eIF1 overexpression, which has been shown to enhance translation initiation stringency ^40,51,52^. Moreover, because eIF1 and eIF5 compete for 40S binding ^42^, this nuclear release of eIF1 is predicted to decrease the fraction of eIF5 bound to the 40S ribosome (Fig. 3C and 3D). To test the role of programmed nuclear release of eIF1 in controlling mitotic translational stringency, we sought to precisely alter the nuclear-cytoplasmic concentrations of eIF1 and eIF5. We first ectopically expressed a nuclear-specific version of eIF5 through the addition of a nuclear localization sequence (NLS-eIF5, Fig. 4A). The addition of this nuclear population of NLS-eIF5 is predicted to reduce translational stringency during mitosis by competing with eIF1 for 40S binding without altering translation in interphase cells by virtue of its nuclear sequestration. Indeed, we found that nuclear-specific eIF5 expression partially suppressed the change in translation stringency during mitosis based on luciferase reporters (3.9-fold to 1.7-fold repression; Fig. 4B). This suggests that creating a nuclear pool of eIF5 is sufficient to partially neutralize the effect of nuclear eIF1 release during mitosis.

Prior work found that eIF1 protein abundance is autoregulated through a negative feedback loop in which eIF1 represses its own translation to maintain homeostatic eIF1 protein levels ^37^ (Fig. 4C). To achieve this, eIF1 harbors an evolutionally-conserved weak translation initiation site such that overexpression of eIF1 increases translation initiation stringency thereby repressing new eIF1 translation ^37^. Consistent with eIF1 autoregulation (Fig. 4C), we found that ectopic expression of GFP-tagged eIF1 under the control of a tetracycline-inducible promoter and an optimal Kozak context resulted in the repression of endogenous eIF1 levels by ∼10-fold during both interphase and mitosis (Fig. 4D; Extended data Fig. 7A-C). In contrast, our model suggests that ectopic expression of nuclear specific eIF1 (NLS-GFP-eIF1) should not enhance translation initiation stringency during interphase, because it is sequestered in the nucleus, but should increase translation initiation stringency during mitosis when it is released from the nucleus. Indeed, ectopic expression of nuclear-specific NLS-GFP-eIF1 repressed endogenous eIF1 levels ∼10-fold during mitosis but had little effect on endogenous eIF1 expression during interphase (Fig. 4D; Extended data Fig. 7A-C). To test whether these effects on endogenous eIF1 levels occur through translational control, we used luciferase reporters harboring the eIF1 5’ UTR. Our analysis indicated that ectopic expression of NLS-GFP-eIF1 preferentially repressed eIF1 translation during mitosis (Fig. 4E). These findings demonstrate that the nuclear sequestration of initiation factors and their release during mitosis can enable switch-like changes to translation during mitosis.

## Nuclear release of eIF1 increases stringency of start codon selection during mitosis

In contrast to the production of additional nuclear eIF1 using the NLS-eIF1 fusion (Fig. 4D), our model predicts that selectively removing the pool of nuclear-localized eIF1 would dampen the differences in translation initiation stringency observed between interphase and mitosis. To test this, we tagged eIF1 at its endogenous locus with either GFP alone (as a control) or GFP and a nuclear export signal (NES) to eliminate the nuclear pool of eIF1 (Fig. 4F; Extended data Fig. 7D-F). In the absence of autoregulation, the addition of a NES to eIF1 would be predicted to display similar total cellular levels of eIF1, but increase the cytoplasmic levels of eIF1 by moving the nuclear eIF1 fraction to the cytoplasm (Extended data Fig. 7E). However, due to the autoregulatory circuit, the increase in cytoplasmic eIF1 levels results in increased translation initiation stringency, which in turn causes the repression eIF1 translation to maintain similar cytoplasmic eIF1 levels in both control GFP-eIF1 and NES-GFP-eIF1 cells (Extended data Fig. 7E). Therefore, NES-GFP-eIF1 cells will have decreased nuclear eIF1 and thus total lower total eIF1 level of eIF1 (Extended data Fig. 7E). Consistent with autoregulation and this model, we found that interphase NES-GFP-eIF1 cells have similar levels of eIF1 in the cytoplasm and decreased levels of nuclear eIF1, with less total eIF1 as compared to GFP-eIF1 cells (Extended data Fig. 7F and 7G).

To directly test the role of nuclear eIF1 release on translational stringency, we measured CUG/AUG translation using luciferase reporters in cells with endogenous GFP-eIF1 and NES-GFP-eIF1 fusions. In GFP-eIF1 cells, the nuclear release of eIF1 increases functional eIF1 during mitosis such that a CUG luciferase reporter is preferentially repressed 2.7-fold during mitosis (Fig. 4G). In contrast, endogenous NES-GFP-eIF1 cells that lack this nuclear release only repressed the CUG reporter 1.2-fold in mitosis relative to the AUG reporter (Fig. 4G; Extended data Fig. 7H). These results indicate that the nuclear release of eIF1 during mitosis accounts for the majority of increase in stringency during mitosis. Together, these results demonstrate that modulating the nuclear/cytoplasmic ratio of eIF1 can differentially regulate translation initiation stringency during interphase and mitosis.

## Depleting nuclear eIF1 sensitizes cells to anti-mitotic chemotherapeutics

To test the biological contributions of increased mitotic translational stringency, we evaluated the functional consequence in cells lacking nuclear eIF1 (endogenous NES-GFP-eIF1), which display a similar interphase translational stringency but decreased mitotic translational stringency compared to control GFP-eIF1 cells (Fig. 4G). We found that endogenously-tagged GFP-eIF1 and NES-GFP-eIF1 HeLa cells proliferated normally and did not display differences in growth under unperturbed conditions (Fig. 5A). This lack of a growth phenotype in unperturbed cells was not surprising as mitosis occurs in ∼60 minutes such that changes in translational regulation over this short timespan will have more modest effects on protein levels, particularly for longer-lived proteins. However, when chromosome segregation is perturbed, either due to physiological damage or following treatment with anti-mitotic chemotherapeutics ^53^, mammalian cells can pause in mitosis for hours to days to enable error correction and safeguard genome stability ^7^. We hypothesized that altered mitotic translational control may become particularly critical under conditions where cells undergo an extended mitotic delay. Indeed, based on propidium iodide staining ^54^ and PARP1 cleavage ^55^, NES-GFP-eIF1 cells lacking nuclear eIF1 were hypersensitive to treatment with high doses of the chemotherapeutic taxol with 75% of NES-GFP-eIF1 cells undergoing cell death in the presence of taxol after 24 hours compared to only 7.5% of GFP-eIF1 cells (Fig. 5B; Extended data Fig. 8A). We observed a similar effect with diverse drugs that cause a mitotic arrest, such as nocodazole or STLC, but not with drugs that do not induce a mitotic arrest (Extended data Fig. 8B). Similarly, following shorter mitotic arrest durations, GFP-eIF1 control cells were able to progress into interphase after release from an 8-hour mitotic arrest, with only 7% undergoing cell death, whereas 35% of NES-GFP-eIF1 cells died following release from a similar 8-hour mitotic arrest (Fig. 5C; Extended data Fig. 8C; Movie S1). Importantly, we were able to rescue both the change in translation initiation stringency and the mitotic cell death by the ectopic expression of NLS-GFP-eIF1 in cells lacking nuclear eIF1 (endogenously-tagged NES-GFP-eIF1; Extended data Fig. 8D). These results suggest that a nuclear pool of eIF1 and increased mitotic translation initiation stringency are required to maintain viability during a mitotic arrest.

We next sought to determine the mechanisms that underlie this mitotic cell death. The depletion of nuclear eIF1 by endogenously tagging eIF1 with NES-GFP specifically reduces translation stringency in mitosis (Fig. 4F-G; Extended data Fig. 7F-H), which is predicted to reduce the relative translation of highly translated mRNAs and increase the relative translation of lowly translated mRNAs (Fig. 2). To test which functional processes would be most affected by this change, we performed gene set enrichment analysis on genes that are differentially translated during mitosis (Extended data Fig. 8E and 8F). We found that nuclear-encoded mitochondrial transcripts were highly translated during mitosis in control cells (Extended data Fig. 8E and 8F). Thus, depleting the nuclear pool of eIF1 is predicted to reduce the relative translation of mitochondrial components and thus mitochondrial fitness specifically during mitosis. To test this, we measured mitochondrial membrane potential as a readout of mitochondrial fitness using the dye tetramethylrhodamine ethyl ester (TMRE) (Fig. 5D-E; Extended data Fig. 8G-H). During interphase, mitochondrial membrane potential was similar in the presence or absence of nuclear eIF1 (Fig. 5D-E; Extended data Fig. 8H), consistent with autoregulation of eIF1 not altering interphase cell physiology. In contrast, during mitosis mitochondrial fitness was significantly reduced in cells lacking nuclear eIF1 (Fig. 5D-E; Extended data Fig. 8H). As decreased mitochondrial activity can lead to cell death ^56^, this may contribute to the mitosis-specific death that occurs in cells lacking nuclear eIF1.

Finally, we analyzed cell cycle progression in cells lacking nuclear eIF1. Mitotically-arrested cells display two mutually exclusive fates—either undergoing cell death in mitosis or slipping out of mitosis into G1 without segregating their chromosomes ^57^. Mitotic slippage results in tetraploid G1 cells and is thought to be a mechanism by which cells develop chemoresistance to avoid anti-mitotic mediated death ^57,58^. Using live imaging, we observed that 60% of GFP-eIF1 cells were able to slip out of mitosis into interphase. In contrast, only 15% of NES-GFP-eIF1 cells exited mitosis in the presence of anti-mitotic drugs (Fig. 5F; Extended data Fig. 8I, Movie S2).

Our recent work demonstrated that one key regulator of mitotic slippage is the ratio of alternative translational isoforms for Cdc20 ^59^, the downstream target of the spindle assembly checkpoint. Leaky ribosome scanning downstream of the first AUG (M1), which has a weak Kozak context, results in the usage of a downstream AUG (M43) to generate a truncated CDC20 protein product. These M1 and M43 Cdc20 protein isoforms display differential inhibition by the spindle assembly checkpoint, with the relative ratios of these isoforms controlling mitotic arrest duration with decreased levels of the truncated Cdc20 isoform strongly delaying mitotic exit ^59^. Based on luciferase mRNA reporters, we found that the translation of CDC20 M1 is decreased during mitosis promoting M43 isoform expression (Extended data Fig. 8J), consistent with eIF1 directly repressing weak Kozak context start sites. In addition, based on western blotting, control GFP-eIF1 cells displayed increased CDC20 M43 translation and steady state protein levels during mitosis (Fig. 5G). In contrast, NES-GFP-eIF1 cells lacking nuclear eIF1 failed to accumulate CDC20 M43 (Fig. 5G). The decreased expression of CDC20 M43 isoform in cells lacking nuclear eIF1 may act as one mechanism to inhibit mitotic slippage in these cells.

Together, our results suggest that the increased stringency of start codon selection during mitosis is mediated by the nuclear release of eIF1. This nuclear release regulates the balance between mitotic slippage and cell death such that modulating nuclear eIF1 levels results in altered sensitivity to anti-mitotic chemotherapeutics (Fig. 6).

## Discussion

Global changes in translation initiation stringency have the potential to substantially alter the proteome. However, such changes have been observed primarily following experimental perturbations that alter ribosome queueing ^41,60^ or the activity or expression of translation initiation factors ^37,39,40,43,52,61^. Physiological conditions in which translation initiation stringency changes remain limited. Here, we find that mitotic cells increase their stringency of start codon selection affecting the relative synthesis of thousands of proteins, including uORFs and alternative translational protein isoforms. Our studies additionally reveal that the nuclear release of eIF1 is a key mechanism for the regulation of mitotic translational stringency. This regulatory control may synergize with prior mechanisms regulating mitotic translation such as the phosphorylation of DENR during mammalian mitosis ^62^, phosphorylation of eIF5A ^63^, and degradation of Ded1p during Yeast meiosis ^64^.

Our work also provides a resource of thousands of alternative translational N-terminal protein isoforms, uORF peptides, and alternative open reading frame whose translation efficiencies vary across the cell cycle. Multiple alternative translational isoforms with both previously-established functions and unknown roles are differentially translated during mitosis (Table S1). The differential synthesis of hundreds of translational isoforms between mitosis and interphase has the potential to impact a wide range of cellular processes, such as we demonstrated for mitochondrial function and cell cycle progression (Fig. 5).

Prior work suggested that cap-dependent translation is reduced during mitosis ^20,21,65,66^. Our results confirm that there is a modest reduction in global translation during mitosis. However, our results also reveal a more targeted change in translation start codon selection during mitosis that influences the specific proteins that are produced. This increase in translational stringency may be important in regulating relative protein levels and alternative translational isoforms, such as for MDM2 (Fig. 1H-I) or CDC20 (Fig. 5G). This translational rewiring may also prevent aberrant translation of RNAs that are released into the cytoplasm during mitosis following nuclear envelope breakdown, including nuclear non-coding RNAs or unspliced RNAs. As the translation of these nuclear RNAs could result in the production of aberrant, toxic proteins, enhancing initiation stringency may help protect the cells from translating these harmful products.

Although the nuclear release of eIF1 and increased translational stringency during mitosis are dispensable for viability in unperturbed HeLa cells, we found that cells lacking nuclear eIF1 are hypersensitive to anti-mitotic chemotherapeutics. Because the median protein half-life is 37.8 hours in HeLa cells ^67^ and 9 hours in H1299 cells ^68^, most proteins may not be depleted dramatically during an unperturbed mitosis, which typically lasts for ∼60 minutes. However, for the 10% of the proteome that displays short half-lives, under conditions where cells experience a mitotic delay, the change in translation stringency would become increasingly important to alter proteomic control and ensure cell survival. For example, oocytes will stay in meiosis II for several days without a nucleus as part of a physiological arrest before fertilization occurs, where altered translational control has the potential to help maintain oocyte viability. In addition, anti-mitotic drugs, such as paclitaxel (taxol), other taxanes, and vinca alkaloids, are widely used as frontline chemotherapeutics for the treatment of a broad range of cancers and act in part by inducing a mitotic delay ^53^. In the absence of nuclear eIF1 release, we found that cells become hypersensitive to taxol, resulting in increased mitotic cell death. Although cancer cells eventually die in the presence of a prolonged mitotic arrest, some evade cell death by “slipping” out of mitosis into a tetraploid G1 state. After undergoing mitotic slippage, some cells are able to maintain their proliferative potential ^59,69,70^. Cells without nuclear eIF1 display both increased cell death during mitosis and a reduced proportion of cells that slip out of mitosis. Thus, our data suggests that depleting nuclear eIF1 could enhance the effectiveness of anti-mitotic cancer drugs and that mitotic cells may be hypersensitive to eIF1 inhibitors.

## Acknowledgments

We thank the members of the Cheeseman, Bartel and Lourido labs, in particular Mary-Jane Tsang, Ekaterina Khalizeva, Yi Fei Tao, Alexandra Nguyen, Arash Latifkar, Chris Guiliano, and Alice Herneisen for helpful discussions; Ekaterina Khalizeva, Eric Smith, Sheamin Khyeam, and Yi Fei Tao for comments on this manuscript; Sarah Cady and Ekaterina Khalizeva for technical assistance; the Whitehead Genome Technology Core for sequencing and the Whitehead Quantitative Proteomics Core for mass spectrometry on the Orbitrap Eclipse.

## Author contributions

Conceptualization: JL, IMC, KX, and DPB

Methodology: JL with support from KX for ribosome profiling and KCS for live cell imaging

Investigation: JL with support from KX for ribosome profiling and GBS for whole cell proteomics, KCS performed live cell imaging mitotic death/slippage, TMRE imaging, and CDC20/PARP1 immunoblots

Formal analysis: JL, KX for linear regression model, and KCS for image quantification

Writing: JL and IMC wrote the manuscript with input from KX, and DPB

Supervision: IMC and DPB

Funding acquisition: IMC, DPB, and JL

## Data and materials availability

Sequencing data and associated analyses were deposited in Gene Expression Omnibus (GSE230189), and the deposition of the mass spectrometry data and plasmids is in progress.

## Methods

### Tissue culture

HeLa, RPE1, HEK293T, NIH3T3, and RAW264.7 cells were cultured in DMEM with 10% heat-inactivated fetal bovine serum, 2 mM L-glutamine and 100 U/mL penicillin-streptomycin at 37°C with 5% CO2.

### Mouse husbandry, oocyte collection, and maturation

The C57BL/6J inbred mouse line was used in this study. Mice were housed in a 12-12 h light/dark cycle with constant temperature and food. The Massachusetts Institute of Technology Department of Comparative Medicine provided daily cage maintenance and health checks. Animal experiments performed in this study were approved by the Massachusetts Institute of Technology Committee on Animal Care (0820-020-23).

For the collection of prophase I-arrested (GV) oocytes, 5- to 8-week-old female mice were super ovulated by intraperitoneal injection of 5–10 I.U. of pregnant mare’s serum gonadotropin (PMSG) (Thermo Fisher, NC1663485). At 48 h post injection, the mice were euthanized, and the ovaries were dissected into EmbryoMax Advanced KSOM Embryo Medium (Millipore Sigma, MR-101-D) supplemented 2.5 μM milrinone (Sigma Aldrich, M4659). Cumulus cells were removed from cumulus-oocyte-complexes by repeated aspiration through a glass pipette then fixed for immunofluorescence. For maturation to meiosis I, GV oocytes were collected as described but using media without milrinone. Denuded oocytes were cultured in Advanced KSOM media at 37°C with 5% CO2 for 8 hours and oocytes where the nuclear envelope has dissolved were fixed for immunofluorescence. For the collection of metaphase II (MII) eggs, 5- to 8-week-old female mice were injected with 5–10 I.U. of PMSG. At 48 h after PMSG injection, the animals were injected with 5–10 I.U. of Human chorionic gonadotropin (hCG) (Millipore Sigma, CG5-1VL). At 16 h post hCG injection, cumulous-enclosed egg complexes were collected from the oviduct into Advanced KSOM media with hyaluronidase (3 mg/ml, Sigma Aldrich, H4272) for 2 min to remove cumulus cells, then fixed for immunofluorescence.

### Cell synchronization

For interphase cells, the mitotic cells were removed by physical disruption (mitotic shake off) and the remaining cells adhered to the plate were harvested. For mitotically arrested cells, HeLa cells were grown to 40% confluency and treated with 2mM thymidine for 24 hours. Cells were released from thymidine arrest by washing twice with warm DMEM and cultured in 8 µM STLC for 16 hours. Mitotically arrested cells were shaken off the plate and harvested. For cycling mitotic cells, HeLa cells were grown to 20% confluency and treated for 2 mM thymidine for 16 hours. Cells were then washed twice with warm DMEM, incubated at 37°C for 8 hours then 2 mM thymidine was added for another 16 hours. After washing twice with warm DMEM and incubation at 37°C for 8 hours, the cycling mitotic cells were shaken off and harvested. G1 cells were isolated by treating mitotically arrested cells with 3 µM AZ3146 (Mps1 inhibitor) for 2 hours and then harvested. G2 cells were arrested using 6 µM Ro-3306 (CDK1 inhibitor) for 24 hours. Ro-3306 arrested cells were washed twice with warm DMEM then treated with STLC or taxol for 2 hours before transfecting for luciferase assays (Extended data Fig. 2C). For mitotically-arrested RAW264.7 cells, cells were grown to 20% confluency then treated with 2 mM thymidine for 16 hours. Cells were released from the thymidine arrest by washing twice with warm DMEM, allowed to grow for 8 hours, and then treated with 2 mM thymidine for 20 hours. Following the second thymidine arrest, cells were washed twice with warm DMEM and allowed to grow in DMEM supplemented with 8 µM STLC for 16 hours. Mitotically-arrested cells were shaken off the plate and used for luciferase assays.

For experiments that required transgene induction and cell synchronization, 1 µg/mL dox was added with the first thymidine synchronization and cells were cultured in the 1 µg/mL dox throughout the experiment.

For CDC20 western blotting experiments, cells were arrested in 2mM thymidine for 23 hours before release into 10 µM STLC. For the “0 hour” mitotic timepoint, mitotic cells were harvested 8 hours after thymidine washout by shake off. For “8 hour” mitotic timepoint, cells were harvested 16 hours after washout by shake off. For PARP1 cleavage western blots, cells were synchronized as above for 8-hour CDC20 but incubated for another 8 hours after mitotic cell shake off (for a total of 16 hour in mitosis).

### Preparation of ribosome profiling and matched RNA sequencing libraries

Interphase, cycling mitotic cells, and STLC-arrested mitotic cells were grown to ∼80% confluency. For elongating ribosome profiling, cells were treated with 100 μg/mL cycloheximide for 2 minutes then harvested. For initiation site sequencing, cells were treated with 2 μg/mL harringtonine for 2 minutes at 37°C, followed by 12.5 μM puromycin for 2 minutes at 37°C then 100 μg/mL cycloheximide and immediately collected. To collect interphase cells, media containing translation inhibitor was poured into a 50mL tube, cells were washed with PBS + 100 μg/mL cycloheximide and then trypsinized in the presence of 100 μg/mL cycloheximide for 3 minutes at 37°C. Cells were then collected using the cold media with inhibitor. For mitotic cells, after addition of cycloheximide, mitotic cells were harvested by shake off and incubated on ice. Cells were centrifuged at 500x g for 3 minutes, washed once with 20 mL PBS+cycloheximide, and the pellet was moved into a new 1.5 mL tube with 1 mL PBS+cycloheximide. Cells were lysed in 900 μL polysome lysis buffer (20 mM Tris pH 7.4, 100 mM KCl, 5 mM MgCl2, 1% (v/v) Triton X-100, 100 μg/mL cycloheximide, 500 U/mL RNaseIn Plus, 1x cOmplete protease inhibitor cocktail) and incubated on ice for 10 minutes. To ensure efficient lysis, cells were passed through a syringe 5 times. Cell debris and nucleus was depleted by spinning the lysate at 1300x g for 10 minutes. 30 μL of the cleared lysate was added to 400 μL TRI Reagent (Invitrogen, AM9738) for input RNA sequencing library. Three 300μL aliquots were generated with the rest of the cleared lysate, flash frozen in liquid nitrogen, and stored at -80°C.

For one aliquot, RNase I (Ambion, AM2294) was added to the lysate, using 40 units per OD260. Samples were incubated at 23° C for 30 minutes before loading onto 10-50% sucrose gradients. Gradients were centrifuged at 36000 RPM for 2 hours at 4°C. Samples were fractionated using on a BioComp gradient fractionator and the 80S fraction was collected and concentrated using a 100 kDa concentration (Amicon, UFC810024). Ribosome protected footprints were separated from individual ribosomes by incubating with release buffer on the 100 kDa concentration column for 10 minutes and centrifuged at 3000x g. The resulting eluant was further treated with 1% SDS and 8 U/mL proteinase K (NEB, P8107S) at 42°C for 20 minutes then phenol chloroform extracted and ethanol precipitated.

RNA was pelleted by spinning at 21000x g for 30 minutes and washed with 70% ethanol. The RNA pellet was then resuspended in 1x denaturing gel loading buffer II (Invitrogen, AM8546G). P32 labelled size markers (20mer and 33mer) were spiked into the ribosome protected fragments, denatured at 65°C, then loaded onto a 10% urea gel alongside Decade RNA marker (Invitrogen, AM7778). 20 nt to 33 nt fragments were gel extracted and precipitated with isopropanol. The 3’ end was dephosphorylated by treatment with T4 PNK followed by ligation of the 3’ pre-adenylated adapter (5′AppTCGTATGCCGTCTTCTGCTTGddC3′) using T4 RNA ligase I. Following 3’ dephosphorylation, rRNA was depleted using a cocktail of complementary biotinylated oligos (http://bartellab.wi.mit.edu/protocols.html) and the ligated product was size selected (41-54 nt) as described above. The 5’ end was phosphorylated using gamma-ATP and T4 PNK then the 5’ adapter (5′ GUUCAGAGUUCUACAGUCCGACGAUCNNNNNNNN 3′) was ligated using T4 RNA ligase I and the ligated product (75-88 nt) was size selected. Fully ligated products were reverse transcribed using SuperScript III reverse transcriptase (Invitrogen, 18080044) then barcoded and amplified by PCR using Kapa HiFi polymerase.

For matched input RNA-seq, 0.05 ng spike-in RNA (equal mixture of in vitro transcribed Fluc, Rluc, GFP mRNA) was added to 500 ng total RNA then the rRNA was depleted with the NEBNext rRNA depletion kit V2 (NEB, E7400L) according to the manufacturer’s instructions. The resulting rRNA depleted RNA was fragmented by incubating at 95°C for 20 minutes in 2 mM EDTA, 12 mM Na_2_CO_3_, 88 mM NaHCO_3_ then ethanol precipitated. The fragmented RNA was size selected and sequenced in parallel with the ribosome protected fragments.

Sequencing was performed on an Illumina HiSeq 2500, with 40 bp standard runs. We performed 2 biological replicates for each condition and drug treatment.

### Translation initiation site identification, quantification, and analysis

Barcodes were trimmed using cutadapt (v3.7) ^71^ with the parameters: -m 15 -u 8 -e 0.1 --match-read-wildcards -a TCGTATGCCGTCTTCTGCTTG -O 1. Trimmed reads were then mapped to the genome using STAR (v2.7.1a) ^72^ with the parameters --runMode alignReads -- outFilterMultimapNmax 1 --outReadsUnmapped Fastx --outFilterType BySJout --outSAMattributes All --outSAMtype BAM SortedByCoordinate.

To identify translation initiation sites, we used RiboTISH^15^. We ran RiboTISH quality with the following parameters: --th 0.40 -l 20,38. We next used RiboTISH predict with harringtonine-puromycin and cycloheximide-protected ribosome footprints including the parameter -e, to predict background estimation model, excluding all possible AUG translation initiation sites. Using the background model from the previous RiboTISH predict, to identify and quantify translation initiation sites, including near-cognate sites, we used RiboTISH predict with the following parameters: -s /pathtobackgroundmodel/ --minaalen 3 --alt --seq --aaseq --inframecount. We then filtered initiation sites with a translation initiation site Q-value of ≤0.05 and frame test q value of ≤0.01. To filter out false positive N-terminal truncations due to unperfect harringtonine treatment (some proportion of ribosomes elongating despite the presence of harringtonine), we filtered out translation initiation sites within the first 16 codons of annotated start codon because on reads within the first 16 codons in our harringtonine-puromycin ribosome profiling data was on average greater 0.5x reads in our cycloheximide ribosome profiling data (likely false positives). We then used RiboTISH tisdiff to compile a count table including reads at translation initiation sites in harringtonine-puromycin treated cells and input mRNA-seq counts. Using reads at the translation initiation sites, normalizations and differential expression analysis were performed using DESeq2. In our DESeq2 design, we used the interaction term to normalize initiation site ribosome protected fragments to input RNA-seq counts, as described in RiboTISH.

To calculate initiation efficiency, we used DESeq2’s median of ratios to get normalized counts. We then normalized initiation site ribosome protected footprint counts by length normalized RNA-seq counts. This metric represents the initiation efficiency, the amount of reads at an initiation site normalized by length-normalized mRNA abundance.

To ensure that harringtonine-bound ribosome protected fragments at translation initiation sites quantitatively represent elongating ribosomes (from cycloheximide-treated cells), we correlated initiation sites to elongating ribosome reads. Initiation site counts and elongating ribosome in-frame counts were obtained from RiboTISH predict. In-frame counts were first normalized to the protein length before correlating initiation site reads and elongating ribosome reads.

We used KPlogo^73^ to generate weighted and unweighted sequence logos. To identify flanking nucleotides that contribute to the change in translation initiation between mitosis and interphase, we used the fold change in initiation efficiency between mitosis and interphase from DESeq2 and the nucleotides flanking the start codon, but excluding the start codon. To recreate the mammalian Kozak motif sequence logo, we took nucleotides flanking only annotated ORFs identified in our data and used KPlogo to generate the motif. In this analysis, each site was weighted equally. We used the KPlogo webserver and default settings to generate the logos.

### Translation efficiency calculation

As described in ^30^, for both cycloheximide ribosome profiling and RNA-seq, only reads that uniquely mapped to coding regions of annotated open reading frames (excluding the first 15 codons and the last five codons) were included in translational efficiency calculations. We used htseq-count (0.11.0) ^74^ with the following parameters to quantify reads mapped to the coding sequence: -f bam -t CDS. An expression cutoff of ≥20 reads in each sample was applied for each gene. Using genes that pass the read cutoff, normalizations of ribosome-footprint reads and RNA-seq reads were performed with DESeq2 ^75^ to calculate the relative translational efficiency. Differential translation efficiency analysis between cell cycle stages was performed with DESeq2 using the interaction term to normalize ribosome protected fragments to input RNA-seq counts.

For reanalysis of ^21,32^, we used their mapped reads and calculated translational efficiency as described above with a read cutoff of at least 20. For ^34^, we mapped, quantified reads, and calculated translational efficiency as described above for our data. For ^33^, we took all genes with >2 TPM, normalized ribosome footprint TPM to input mRNA TPM, and then correlated translational efficiencies between conditions.

### Analysis of N-terminal degrons

For analysis of N-terminal degrons, the second amino acid (amino acid immediately downstream of initiator methionine) for each translational isoform was counted for the proportion of arginine, lysine, glycine, and cysteines as these N-terminal amino acids direct N-terminus mediated protein decay ^27^.

### Genome references and gene annotations

Human genomic sequences and annotations were downloaded from the GENCODE website (release 25, GRCh38.p7, primary assembly or main annotation). For translational efficiency calculations, annotations for protein-coding genes were extracted, and the isoform with the longest ORF was selected to represent that gene. For genes with multiple isoforms with the same longest ORF length, the longest transcript was used. The coding sequences of Fluc, Rluc, eGFP (spike-ins) was added to the annotation.

Annotated ORFs reflect human gencode V25 annotations, other ORFs correspond to internal out-of-frame or 3’ UTR ORFs, uORFs are ORFs in the 5’ UTR (including overlapping uORFs), truncations are in-frame of the annotated ORF but start downstream of the annotated start, and extensions are in-frame and upstream of the annotated without an intervening stop codon.

For translational efficiency analysis, we used the MitoCarta3.0 annotation for nuclear-encoded mitochondrial genes ^76^. The annotation from Human Protein Atlas was used for nucleolar genes ^77^.

### Linear regression

For each translation initiation site, we compiled the following features for the linear regression: 1) Upstream 5ʹ-UTR 1–6mer frequency; 2) Downstream exon 1–6mer frequency; 3) Total mRNA GC content and downstream exon (198 nt) GC content; 4) CodonScores, a metric used for modeling the preference of translation initiation codons ^78^; 5) Kozak Similarity Score, a metric used for modeling the similarity between Kozak consensus sequence and the sequence context surrounding translation initiation codons ^79^; 6) Translation initiation codon identity; 7) Translational efficiency during interphase.

All features and the target value (log_2_ fold change of initiation efficiency) were standardized using StandardScaler in the python scikit-learn package. A cutoff of averaged 50 reads between 2 replicates was used to filter translation initiation sites. The remaining initiation sites were stratified into 10 folds based on labels of whether an initiation site is AUG or not, using StratifiedKFold in the python scikit-learn package. We used 9 folds data for training and 1 fold data as held-out for testing and rotated the held-out data for all 10 folds. For each rotation, we trained 5 models using stochastic gradient descent with SGDRegressor in the python scikit-learn with the following parameters: alpha = 0.01, l1_ratio = 0.5, max_iter = 100, penalty = ‘elasticnet’, eta0 = 0.001, power_t = 0.25, n_iter_no_change = 10. The model with the highest Pearson correlation coefficient was chosen for prediction on the held-out data.

For linear modeling of translational efficiency data, we used geom_smooth. In the linear modeling we used mitosis as the independent variable and interphase translational efficiency as the dependent variable (regression of x on y).

### Gene set enrichment analysis

We used the fgsea package in R to perform gene set enrichment analysis^80^. We used a permutation number of 10000. For ontology gene sets, we downloaded GO pathways (c5.go.v7.4.symbols.gmt or c5.go.cc.v7.4.symbols.gmt) from Human MSigDB Collections (UCSD/Broad).

### Propidium iodide live/dead assay

For assays related to mitotic cell viability in the presence of translational inhibitors, HeLa cells were arrested in S phase using 2mM thymidine (Sigma) for 24 hours. Cells were washed twice using warm DMEM and released into DMEM containing 8 μM STLC for 16 hours. Mitotically arrested cells were shake off, replated into a new well and incubated in 100 μg/mL cycloheximide with or without 100 μM and MG132 for various times. Propidium iodide (Invitrogen, P3566) was added to the cell media at 1 μg/mL and incubated at 37C for 15 minutes. Following incubation, cells were strained and the population of propidium iodide positive cells were monitored using the BD FACSymphony A1 Cell Analyzer (BD Biosciences) and analyzed using FlowJo V10.7.1. We first gated cells using forward scatter and side scatter area prior to calculating the proportion propidium iodide positive cells.

For mitotic cell viability (Fig. 5), cells were arrested in S phase using 2mM thymidine (Sigma) for 16 hours. Cells were washed twice using warm DMEM and released into DMEM containing 1mM taxol, 8 μM STLC, or 1 μg/mL nocodazole for 8 hours. Mitotic cells were shaken off, replated into a new well, and incubated for 24 hours prior to propidium iodide staining. 100 μM Sodium arsenite (Sigma, S7400-100G) or 10 μM MG132 (VWR, 89161) was added to cycling asynchronous cells for 24 hours prior to viability measurements (Extended data Fig. 8B). We note that this concentration of MG132 inhibits entry into mitosis so does not result in accumulation of mitotic cells. Propidium iodide staining and flow cytometry was performed as described above.

### In vitro mRNA synthesis

Plasmids containing the T7 promoter and a reporter of interest were amplified by PCR to generate a template for in vitro transcription. The PCR product was purified using EconoSpin TM All-in-1 Mini Spin Columns (Epoch Life Science, 1920-250) and eluted with RNase free water. 1 µg of PCR product was used in an in vitro transcription reaction using the HiScribe T7 High Yield RNA Synthesis kit (NEB, E2040S) in the presence of 20units/mL SUPERaseIn. Free nucleotides and abortive transcripts were removed using P30 columns (Bio Rad, 7326251) and the resulting eluate was purified by phenol chloroform extraction. Purified in vitro transcribed RNA was capped using the Vaccinia Capping System (NEB, M2080S) according the manufacturer’s protocol, passed through a P30 column and purified by phenol chloroform extraction. The resulting capped RNA was polyadenylated using *E. coli* Poly(A) Polymerase (NEB, M0276L), cleaned up using P30 column and phenol chloroform extraction. Capped and polyadenylated RNA was quantified by nanodrop, aliquoted, flash frozen, and stored at -80C.

### mRNA transfections

Cells were grown to ∼80% confluency before transfection in 24 well plates. For mitotic cells, cells were isolated by mitotic shake off and concentrated to ∼80% confluency before transfecting, to match the density of interphase cells. Cells were transfected with Lipofectamine MessengerMAX (Thermo Fisher, LMRNA008), according to manufacturer’s instructions. We transfected 500 ng mRNA per well using 1 µL Lipofectamine MessengerMAX. For luciferase assays, cells were harvested 2.5-3 hours post transfection.

### Plasmid transfection for luciferase assays

To test the activity of tagged eIF1 constructs (Extended data Fig. 5A), we co-transfected AUG or CUG nano luciferase, firefly luciferase, and the indicated overexpression constructs. We used the nano luciferase reporter constructs from ^60^. Cells grown to ∼80% confluency in a 24 well plate were transfected with 37.5 ng AUG or CUG nano luciferase plasmid, 100 ng firefly luciferase plasmid and 112.5 ng GFP or different eIF1 plasmids using Lipofectamine 2000 (Thermo Fisher). After 24 hours, cells were harvested for luciferase assays. For the linker, we used a 50 amino acid glycine-serine linker.

### Luciferase assays

Luciferase assays were performed using the Dual-Luciferase Reporter Assay System (Promega) according to the manufacturer’s protocol. For interphase cells, cells were washed once with PBS and lysed in passive lysis buffer on the plate (Promega). For mitotic cells, cells were isolated by mitotic shake off (to separate them from any contaminating interphase cells on the plate), spun down at 500g x for 3 minutes, washed once with PBS, then lysed in passive lysis buffer (Promega). Luciferase assays were carried out in using the GloMax Discover Microplate Reader (Promega).

### Polysome gradient for mass spectrometry analysis

Lysates and sucrose gradients were prepared as described for ribosome footprint profiling except SUPERaseIn was used instead of RNaseIn Plus. We collected free, 40S, 60S, and 80S ribosome fractions. Each fraction was TCA precipitated using 1/5^th^ (v/v) volume of 100% TCA overnight on ice. TCA precipitant was pelleted by spinning at 21000x g for 30 minutes. The protein pellet was washed with 1 mL -20°C acetone and centrifuged at 21000x g for 15 minutes. The acetone wash was repeated 2 times for a total of three washes. The protein pellet was dried to get rid of residual acetone by speed vac then resuspended in 24 µL 1x S-trap lysis buffer and quantified using the BCA protein assay kit (Pierce). After quantification, samples were either prepared used for mass spectrometry or western blot analysis.

### Immunoprecipitations

Polyclonal GFP antibodies was coupled to Protein A beads as described in ^81^. For GFP-eIF1 IP experiments, 5 15cm plates of cells were dislodged using PBS+ 5 mM EDTA resuspended in DMEM and spun down at 500 x g for 3 minutes. Cell pellet was washed twice in ice cold PBS then resuspended 900uL 20 mM Tris pH 7.4, 100 mM KCl, 5 mM MgCl2, 1% (v/v) Triton X-100, 20units/mL SUPERaseIN, and 1x cOmplete protease inhibitor cocktail then flash frozen in liquid nitrogen and stored at -80°C. Lysates were thawed on ice then sonicated using the Covaris S22, in a 1 mL milliTUBE with AFA fiber for 5 mins with the following settings, PIP: 140, Duty Factor: 5%, CPB: 200, Setpoint temperature: 6°C. Samples were cleared by spinning at 1300x g for 10 minutes, the resulting supernatant was then mixed with GFP-agarose beads, and rotated end-over-end for 1 hour at 4°C. Beads were washed 5x with 20 mM Tris pH 7.4, 100 mM KCl, 5 mM MgCl2, 1% (v/v) Triton X-100, 10 μg/mL leupeptin/pepstatin/chymostatin, and 20units/mL SUPERaseIN for 5 minutes at 4° C while rotating end-over-end. Bound protein was eluted with 100 mM glycine pH 2.6, precipitated by addition of 1/5th volume TCA at 4°C overnight. TCA precipitant was washed 3 times with -20°C acetone and dried using a speed vac.

For endogenous eIF1 IP experiments, the same protocol for GFP-eIF1 IP was used except we used α-eIF1 antibody couple to protein A beads.

### Recombinant protein expression affinity purification of eIF1 antibody

BL21(DE3) carrying the pRARE tRNA plasmid were transformed with the appropriate plasmid and plated on LB-agar plates containing the appropriate antibiotic. Overnight liquid cultures of LB supplemented with antibiotics and grown overnight at 37C from single colonies. The saturated overnight culture was diluted 1:100 and grown to an OD600nm of 0.6-0.7 at 30C. Cells were shifted to 16C and induced with 0.3 mM isopropyl b-D-1-thiogalactopyranoside and incubated for 16 hr. Cells were collected by centrifugation, resuspended in lysis buffer, and flash frozen in liquid nitrogen. Cell pellets were resuspended in Lysis buffer (1X PBS supplemented with 250 mM NaCl, 0.1% Tween-20, 1 mM dithiothreitol, and 1 mM phenylmethylsulfonyl fluoride). Cells were disrupted by sonication, and the lysate was cleared by centrifugation. The lysate was applied to 0.5 mL of glutathione agarose (Sigma-Aldrich) per liter of culture for 1 hr at 4C. Agarose was washed three times in Lysis Buffer, and proteins were eluted using Elution Buffer (50 mM Tris-HCl, 75 mM KCl, 10 mM reduced glutathione, pH 8.0). Eluted proteins were dialyzed overnight twice in PBS then in PBS+50% glycerol overnight. Proteins were flash frozen in liquid nitrogen, and stored at -80°C.

### Affinity purification of eIF1 antibody

Purified GST-eIF1 was sent to Covance for antibody production in rabbit. The serum from immunized rabbit was depleted for antibodies against GST by circulating the serum over GST coupled to a 5 mL HiTrap-NHS column (Cytiva, 17071701). To ensure complete depletion of GST antibodies, the resulting flowthrough was circulated through a fresh GST column again. Following this anti-GST depletion, eIF1 antibody was affinity purified by circulating the 2x GST depleted serum through a GST-eIF1 coupled 1 mL HiTrap-NHS column (Cytiva, 17071601). The affinity purified antibody was dialyzed in PBS+50% glycerol, flash frozen in liquid nitrogen, and stored at -80°C. We note that for unknown reasons, untagged eIF1 could not be coupled to any column, which is why we depleted GST antibodies followed by purified eIF1 antibodies using this protocol.

### Whole cell quantitative proteomics

One 15 cm plate of interphase or mitotically arrested cells was isolated as described above. Cells were washed twice in PBS then lysed in RIPA buffer (150 mM NaCl, 50mM Tris pH 7.4, 1% NP-40, 0.5% sodium deoxycolate, and 0.1% SDS) on ice. Lysate was sonicated, cleared by centrifugation, and quantified by BCA protein assay. Lysate was prepared for mass spectrometry using the S-trap protocol described below.

### Mass spectrometry sample preparation

Proteins were digested and cleaned up using a modified version of the S-trap protocol (Protifi, V4.7). TCA precipitant was resuspended in 1x lysis buffer (5% SDS, 50 mM TEAB pH 8.5), denatured at 95°C for 10 minutes in the presence of 20 mM DTT. Samples were alkylated with 40 mM iodoacetamide for 30 minutes at room temperature then acidified to a final concentration of 2.5% v/v phosphoric acid. 6X volume of S-trap binding/wash buffer was added then loaded onto S-trap mini columns. Samples were spun at 4000x g for 30 seconds and washed 4 times with 150 µL S-trap binding/wash buffer by spinning at 4000x g for 30 seconds. After the last wash, the column was dried by spinning at 4000x g for 1 minute. Proteins on column were digested overnight at 37°C in a humidified incubator with 20 µl of 50 mM TEAB pH 8.5 containing 1 µg of trypsin. Peptides were eluted using 40 µl of 50 mM TEAB, 0.2% formic acid then 50% acetonitrile/0.2% formic acid. The eluted peptides were quantified using the Themo fluorescent peptide quantification kit, flash frozen then lyophilized.

### TMT labelling and peptide fractionation

1.5 µg trypsinized peptides were dissolved in 50 mM TEAB pH 8.5 and labelled using the TMT10plex Isobaric Labeling Reagent Set (Thermo Fisher Scientific, 90111). Each sample was labeled with TMT10plex reagents at a 20:1 label:peptide w/w ratio, for 1 hour at room temperature. TMT labeling reaction was quenched with 0.2% hydroxylamine for 15 minutes at room temperature. The samples were pooled on ice, flash frozen and lyophilized. Pooled TMT-labelled peptides were cleaned and fractionated using the Pierce High pH Reversed-Phase Peptide Fractionation Kit (Thermo Fisher Scientific, 84868) according to manufacturer’s instruction for TMT experiments. After fractionation, samples were flash frozen and lyophilized.

For quantitative polysome profiling experiments, we performed 2 separate TMT10plex experiments. Each experiment contained interphase and mitotic, free, 40S, 60S, and 80S peptides. The two replicates were processed and analyzed independently.

For quantitative eIF1 interaction experiments, a single TMT experiment included 2 biological replicate GFP immunoprecipitations from GFP, GFP-eIF1, and NLS-GFP-eIF1 expressing cells.

### Mass spectrometry data acquisition

Lyophilized peptides were resuspended in 0.1% formic acid to a final concentration of 250 ng/uL. Mass spectrometry was performed using an Exploris 480 Orbitrap mass spectrometer equipped with a FAIMS Pro source connected to an EASY-nLC chromatography system. Our samples are separated using either a 15 or 25 cm analytical column (PepMap RSLC C18 3 µm, 100A, 75 µm). For all mass spectrometry experiments, peptides were separated at 300 nl/min on a gradient of 6–21% B for 41 min, 21–36% B for 20 min, 36–50% B for 10 min, 50–100% B over 15 min, 100–2% B for 6 minutes, and then 2–100% for 6 minutes. The orbitrap and FAIMS were operated in positive ion mode with a positive ion voltage of 1800 V; with an ion transfer tube temperature of 270°C; using standard FAIMS resolution and compensation voltages of –50 and –65 V (injection 1) or –40 and –60 (injection 2). Full scan spectra were acquired in profile mode at a resolution of 120,000, with a scan range of 350– 1200 m/z, automatically determined maximum fill time, standard AGC target, intensity threshold of 5×10^3^, 2–5 charge state, and dynamic exclusion of 30 seconds.

For mass spectrometry experiments related to endogenous eIF1 IP-MS (Extended data Fig. 6C), mass spectrometry was performed using an Eclipse Orbitrap mass spectrometer equipped with a FAIMS Pro source connected to an Vanquish Neo nLC chromatography system. Lyophilized peptides were resuspended in 0.1% formic acid to a final concentration of 250 ng/uL. Our samples are separated using a 25 cm analytical column (PepMap RSLC C18 3 µm, 100A, 75 µm). For all mass spectrometry experiments, peptides were separated at 300 nl/min on a gradient of 3–25% B for 57 min, 25–40% B for 17 min, 40–95% B for 10 min, 95% B over 6min, and then an equilibration of the column to 3% B at the end of the run using 0.1% FA in water for A and 0.1% FA in 80% acetonitrile for B. The orbitrap and FAIMS were operated in positive ion mode with a positive ion voltage of 2100 V; with an ion transfer tube temperature of 305°C and a 4.2 l/min carrier gas flow, using standard FAIMS resolution and compensation voltages of –50 and –65 V. Full scan spectra were acquired in profile mode at a resolution of 120,000 (MS1) and 50,000 (MS2), with a scan range of 400–1400 m/z, custom maximum fill time (200ms), custom AGC target (300% MS1, 250% MS2), isolation windows of m/z 0.7, intensity threshold of 2×104, 2–6 charge state, dynamic exclusion of 60 seconds, and 38% HCD collision energy.

### Mass spectrometry analysis

Raw files were analyzed in Proteome Discoverer 2.4 (Thermo Fisher Scientific) to generate protein and peptide IDs using Sequest HT (Thermo Fisher Scientific) and the Homo sapiens protein database (UP000005640) with GFP. The maximum missed cleavage sites for trypsin was limited to 2. Precursor and fragment mass tolerances were 10 ppm and 0.02 Da, respectively. The following post-translational modifications: dynamic phosphorylation (+79.966 Da; S, T, Y), dynamic oxidation (+15.995 Da; M), dynamic acetylation (+42.011 Da; N-terminus), dynamic Met-loss (–131.04 Da; M N-terminus), dynamic Met-loss+acetylation (–89.03 Da; M N-terminus), and static carbamidomethyl (+57.021 Da; C). For TMT experiments, we included static TMT6plex (+229.163 Da; any N-terminus), and static TMT6plex (+229.163 Da; K). TMT 10plex isotope correction values were accounted for (Thermo Fisher; 90111 LOT# VK306786). Peptides identified in each sample were filtered by Percolator to achieve a maximum FDR of 0.01 ^82^. For the quantitative TMT polysome proteomics, whole cell proteomics, and endogenous eIF1 IP experiments, protein abundances were normalized on the total protein amount. For any quantitative GFP-IP experiments, protein abundances were normalized to GFP to account for differences in IP efficiency.

### Immunoblots

Cell or TCA precipitated protein pellets were resuspended in 1x Laemmli sample buffer (100 mM Tris pH6.8, 12.5% glycerol (v/v), 1% SDS (w/v), 0.1% bromophenol blue (w/v), 200 mM β-mercaptoethanol) then boiled at 95°C for 5 minutes. Whole cell extracts were sonicated at 10% amplitude for 5 seconds using the Branson Digital Sonifier 450 Cell disrupter before boiling to sheer genomic DNA. Samples were separated by SDS-PAGE and transferred to PVDF (VWR). Blots were rinsed once with TBST then blocked in 5% milk at room temperature for 1 hour. Primary antibodies were diluted in 5% milk and incubated with the blot overnight at 4°C, washed for 5 minutes with TBST 4x, incubated with secondary antibody in 5% milk for 1 hour at room temperature, followed by another 4 washed with TBST, and rinsed with PBS twice. For chemiluminescence (CDC20 and PARP1 immunoblot experiments, clarity enhanced chemiluminescence substrate (Bio-Rad) was added to the membrane according to the manufacturer’s instructions. Membrane was imaged with a KwikQuant Imager (Kindle Biosciences). For LI-COR, blots were imaged using an Odyssey Clx machine (LI-COR) and analyzed with the Image Studio software (LI-COR).

The following antibodies were used: anti-eIF2α (1:1000, Cell Signaling Technology, #9722), anti-phospho-eIF2α Ser51 (1:1000, Cell Signaling Technology, #3398), anti-eIF1 (1:1000, Cell Signaling Technology, #12496), anti-eIF1 (1ug/mL, this paper), anti-eIF5 (1:2000, Proteintech, 11155-1-AP), anti-RPS3 (1:1000, Cell Signaling Technology, #9538), anti phospho-histone H3 (Ser10) (1:1000, Cell Signaling Technology, #9701), anti-CDC20 (1:200, Santa Cruz Biotechnology, sc-13162), anti-PARP (1:1000, Cell Signaling Technology, #9532), anti-GAPDH (1:4000, Santa Cruz Biotechnology, sc-47724) and, anti-alpha tubulin (1:4000, Sigma-Aldrich, T9026). The following secondary antibodies were used: IRDye 680RD Goat anti-Rabbit (LI-COR 92668071), IRDye 680RD Goat anti-Mouse (LI-COR 92668070), IRDye 800CW Goat anti-Rabbit (LI-COR 92632211), IRDye 800CW Goat anti-Mouse (LI-COR 92632210). For Chemiluminescence, we used HRP-conjugated secondary antibodies (Kindle Biosciences, R1005 or R1006).

### Cell cycle analysis by flow cytometry

Cells were synchronized as described above in a 6 well plate. Interphase cells were trypsinized, washed once in PBS, and pelleted. Mitotic cells were isolated by shake off, spun down, and washed once in PBS. To fix cells, the cell pellet was resuspended in 500 µL PBS then 9.5 mL -20°C ethanol was added to the cell drop wise while lightly vortexing, and incubated on ice for >30 minutes. Following fixation, cells were washed once in PBS+0.3% BSA (w/v), then once in PBS + 3% BSA (w/v) + 0.1% Triton X-100 (v/v), and then blocked using antibody dilution buffer (AbDil, 20 mM Tris-HCl, 150 mM NaCl, 0.1% Triton X-100, 3% bovine serum albumin, 0.1% NaN_3_, pH 7.5) on ice for 30 minutes. After blocking, cells were resuspended in 1:1000 anti-phospho-histone H3 Ser10 (Abcam, ab5176) diluted in AbDil and incubated overnight at 4°C, rotating end over end. Cells were then washed once with PBS + 3% BSA + 0.1% Triton X-100, and incubated in 1:300 goat anti-rabbit cy5 (Jackson ImmunoResearch Laboratories) diluted in AbDil for 1 hour on ice. Cells were then washed once with PBS + 3% BSA + 0.1% Triton X-100, and incubated with PBS + 20 μg/mL hoescht for >30 minutes on ice. Cells were then strained and measurements were made using the BD LSRFortessa Cell Analyzer (BD Biosciences). Data was analyzed using FlowJo. We gated for live cells and singlets prior to any quantification.

### Immunofluorescence and quantification

HeLa and NIH3T3 cells were seeded on poly-L-lysine coated coverslips and fixed in PBS + 4% formaldehyde + 0.1% Triton X-100 at room temperature for 15 minutes. Following fixation, cells were washed 3 times with PBS + 0.1% Triton X-100 and blocked in AbDil for 30 minutes - 1 hour. Following blocking, cells were stained at room temperature for 1 hour with α-eIF1 antibody (1 µg/mL) and α -alpha tubulin (1:1000, Sigma-Aldrich, T9026) diluted in AbDil. Cells were washed with PBS + 0.1% Triton X-100, 3 times then incubated in secondary antibody diluted in AbDil for 1 hour at room temperature. After secondary, cells were stained in hoescht for 15 minutes at room temp, then washed 3 times with PBS + 0.1% Triton X-100 before mounting in PPDM (0.5% p-phenylenediamine and 20 mM Tris-Cl, pH 8.8, in 90% glycerol) and sealed with nail polish.

For oocyte and cumulus cell staining, cells were fixed in PBS + 2% formaldehyde + 0.1% Triton X-100 for 20 minutes then washed twice with blocking buffer (PBS + 0.3% BSA + 0.1% Tween-20). Oocytes were then permeabilized in PBS + 0.3% BSA + 0.1% Triton X-100 for 20 minutes followed by 3 consecutive blocking buffer washes for 10 minutes each. Cells were then incubated at room temperature for 1 hour with primary antibody α-eIF1 antibody (1 µg/mL) and α-alpha tubulin (1:1000, Sigma-Aldrich, T9026) diluted in blocking buffer, followed by 3 consecutive 10 minute washes in blocking buffer. Oocytes were stained with secondary antibody was diluted in blocking buffer followed by 3 consecutive 10 minute washes in blocking buffer. Oocytes were mounted in VectaShield with DAPI (Vector Laboratories, H-1200) and sealed with nail polish. Images were taken on the Deltavision Ultra (Cytiva) system using a 60x/1.42NA objective. 8 μm images were taken and 20 μm were taken for cells and oocytes, respectively, with z-sections of 0.2 μm. All images presented were deconvolved and max projected. Background normalized eIF1 signal was quantified using a custom cell profiler pipeline.

### Live cell fluorescence microscopy

Cells on glass bottom plates were incubated in 0.1 μg/mL hoescht for >30 minutes. The GFP transgene was induced by addition of 1 µg/mL dox for 24 hours prior to imaging. Images were taken on the Deltavision Ultra (Cytiva) system using a 60x/1.42NA objective. 8 μm images were taken with z-sections of 0.2 μm. All images presented were deconvolved and max projected.

### Live cell imaging for mitotic death and slippage

Cells were first seeded in 12-well polymer-bottomed plates (Cellvis, P12-1.5P) and moved to CO2-independent media (Gibco) supplemented with 10% FBS, 100 U/mL penicillin and streptomycin, and 2 mM L-glutamine for imaging at 37°C. Phase contrast images were acquired on a Nikon eclipse microscope equipped with a sCMOS camera (ORCA-Fusion BT, Hamamatsu) using a Plan Fluor 20X/0.5 NA objective at 5- or 10-minute intervals. For prolong mitotic arrest experiments, cells were imaged in 1 mM Taxol 1 hour post drug addition for 50 hours. For STLC wash out experiments cells were arrested in S phase using 2mM thymidine (Sigma) for 23-24 hours. Thymidine was removed and cells were further incubated in STLC for 16 hours. STLC was then washed out before imaging 30 minutes post washout in imaging media for 24 hours.

### TMRE staining and mitochondrial fitness quantification

For mitotic studies cells were arrested in 2 mM thymidine for 23 hours before release into 10 µM STLC. 10 hours later mitotic cells were shaken off and incubated before staining with 10 nM TMRE (Invitrogen, T669) for 30 minutes. Cells were then washed and imaged in glass bottom dish (cellvis) after 5 min centrifugation at 200x g in CO2 independent imaging media at t=26h post release using Nikon eclipse at 10x magnification and Cy3 fluorescence filter setup (see time lapse). For interphase asynchronous cells were trypsinized before staining and imaging. As a positive control for decrease in mitochondrial fitness, 50 µM CCCP (Thermo Scientific, 228131000) was added to cells for 15 minutes prior to TMRE staining. TMRE signal was background normalized and quantified using a custom cell profiler.

### Molecular biology and cell line generation

For AUG and CUG luciferase mRNA reporter experiments, the CD9 5’ UTR was used in all constructs ^20^. MDM2 (ENST00000258149.11), CD9 (ENST00000009180.10), eIF1 (ENST00000469257.2), and CDC20 (ENST00000310955.11), 5’ UTR, nano luciferase, EMCV IRES and firefly luciferase were cloned downstream of the T7 promoter using Gibson Assembly. For MDM2 mRNA reporters, the AUG for both uORFs were mutated from AUG to AAG. For eIF1 mRNA reporters, we used Gibson assembly to generate versions with wild-type (uaucguAUGu) or perfect Kozak (gccaccAUGg) contexts. For the CDC20 M43 reporter, a premature stop codon was added downstream of M1 so only M43 initiation is measured by nano luciferase activity. The eIF1 CDS was cloned downstream of GST in pGEX6P1 and used for recombinant protein production.

mEGFP-eIF1, and mEGFP-eIF5 constructs were cloned downstream of the tetracycline-responsive promoter. For eIF1 and eIF5 constructs, a 50 amino acid glycine-serine linker separated mEGFP and eIF1/eIF5. mEGFP-eIF1 constructs also contained a downstream EMCV IRES - mCherry of eIF1. For nuclear specific eIF1, the cMyc NLS (PAAKRVKLD) ^83^ was upstream and the SV40 NLS (PKKKRKV) ^84^ was downstream of the mEGFP. For cytoplasmic specific eIF1, the NES17 (IDELAKALPDLNLD) was upstream and the BIV REV NES (IQQLEDLVRHMSL) ^85^ was downstream of the mEGFP. For nuclear specific eIF5, the nucleoplasmin NLS (KRPAATKKAGQAKKKK) and cMyc NLS was tagged at the N-terminus and the C-terminus of eIF5 was tagged with mEGFP – SV40 NLS (PKKKRKV). Donor plasmids contained puromycin N-acetyltransferase (puromycin resistance), reverse tetracycline-controlled transactivator, and the tetracycline inducible transgene flanked by piggyBac inverted terminal repeats. To generate stable tetracycline inducible cell lines, 1 µg donor plasmid was co-transfected with 400 ng piggyBac transposase plasmid using lipofectamine 2000. 2 days post-transfection, cells were selected with 0.4 µg/mL puromycin for 3 days. For experiments related to rescue of translational stringency and mitotic viability in cells lacking endogenous nuclear eIF1 (Extended data Fig. 8D), the tetracycline-inducible NLS-GFP-eIF1 was transfected into endogenously tagged NES-GFP-eIF1 cells.

For endogenous eIF1 tagging HeLa cells were transfected with 500 ng pX330-eIF1 gRNA plasmid (gRNA sequence 5’ - AGAGTGGAGGTTCTGGATAG - 3’) and 500ng GFP-eIF1 or NES17-mEGFP-BIV REV NES-eIF1 recombination plasmid using lipofectamine 2000. Homology arms of ∼750 bp on either side was used in our recombination template. Fresh media was swapped 16 hours post-transfection. 24 hours after swapping media, the cells were moved into a 15 cm plate and allowed to grow for 72 hours. GFP positive cells were bulk sorted using fluorescence activated cell sorting. After the first sort, cells were grown for another 48 hours to recover. Cells were then subjected to a second round of sorting, where we sorted the top 15% GFP positive cells, to enrich for homozygotes. For monoclonal lines, these GFP positive cells were single cell sorted. Multiple clones were tested and we have shown one representative clone.

To indelibly deplete cytoplasmic eIF1 in order to monitor nuclear eIF1 localization throughout the cell cycle (Extended data Fig. 5H), the GFP nano body (VhhGFP4) was cloned upstream of the SPOP ΔNLS degron ^86^ and IRES2 mCherry. This construct was cloned downstream of the tetracycline-responsive promoter. Donor plasmid also contained the puromycin resistance marker, reverse tetracycline-controlled transactivator, the Vhh-GFP4-SPOPdeltaNLS-IRES2-mCherry transgene, flanked by safe harbor AAVS donor homology arms. Endogenously tagged GFP-eIF1 cells were co-transfected with, 500 ng donor plasmid and 500 ng pX330-sgRNA AAVS1 using lipofectamine 2000. 2 days post-transfection, cells were selected with 0.4 µg/mL puromycin for 3 days.

To generate lentivirus, HEK293T cells at ∼90% confluency in a 6 well plate was transfected with 1.2 µg lentiCas9-sgRNA transfer plasmid, 1 µg psPAX2 packaging plasmid, and 0.4 µg vsFULL envelope plasmid using Xtremegene-9 (Roche). After 16 hours, the media was swapped. The media was collected 24 hours after and stored at -80°C. The following RNA sequences were used: sgEIF1-a: 5’-GTAATTGAGCATCCGGAATA - 3’ and sgEIF1-b: 5’ - AGAGTGGAGGTTCTGGATAG - 3’

### Competitive growth assays

For GFP-eIF1 and NES-GFP-eIF1 competitive growth assays, these cells were competed against HeLa expressing mCherry. The relative GFP:mCherry ratio is displayed. To directly compete GFP-eIF1 and NES-GFP-eIF1, the GFP-eIF1 cells were infected with eBFP2 lentivirus and NES-GFP-eIF1 with mCherry lentivirus. The relative mCherry:eBFP2 ratio is displayed. The relative ratio of fluorescent cells were monitored every 3 days using the BD LSRFortessa Cell Analyzer (BD Biosciences).

---

**Extended data Fig. 1.**
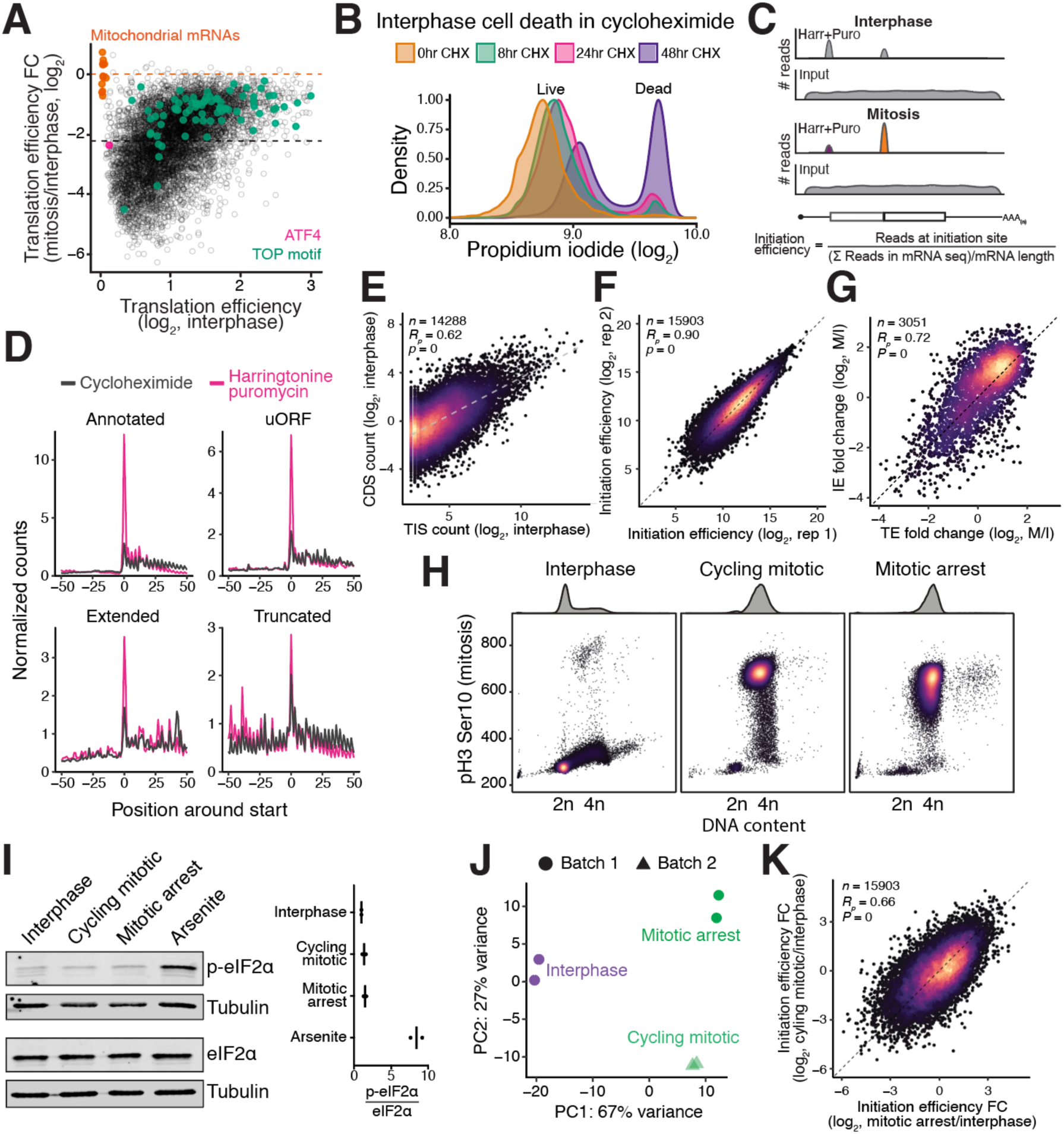
Translation initiation site analysis quantitatively captures translation. (**A**) Translational efficiency fold change between mitotically arrested and interphase cells, normalized to mitochondrial footprints. Black dotted line represents median translational efficiency fold change for cytoplasmic mRNAs between mitotically arrested and interphase cells. Orange line represents median of translational efficiency FC for the 13 mitochondrial coding mRNAs. The median translational efficiency of cytoplasmic mRNAs is 4-fold more repressed than mitochondrial mRNAs in mitosis relative to interphase. (**B**) Distribution of live/dead cells showing the cell viability of asynchronous HeLa cells treated with cycloheximide for various times. (**C**) Normalization strategy to calculate translation initiation efficiency for each start site. Reads mapping to the translation initiation site, quantified by RiboTISH, were normalized to matched length-normalized input RNA seq counts. (**D**) Metagene plot of normalized reads from annotated (top left), uORFs (top right), N-terminal extensions (bottom left), and N-terminal truncations (bottom right) from translation initiation site sequencing (magenta) and elongating ribosome profiling (dark grey). X-axis represents the position around the predicted start codon and Y-axis is the normalized ribosome-protected footprints at each position. (**E**) Translation initiation site counts (Y-axis) correlates with length-normalized elongating ribosome counts (X-axis). The plot is shown for a single representative replicate from interphase cells. (**F**) A scatter plot shows that biological replicates for translation initiation site sequencing are highly correlated and reproducible. (**G**) Scatter plot showing that translation initiation efficiency fold-change and translational efficiency fold-change between mitosis and interphase correlate with each other. Annotated ORFs with only a single identified translation initiation site were included in this analysis. (**H**) Cell cycle analysis of interphase, cycling mitotic, and STLC-arrested mitotic cells indicating efficient synchronization. Cells were stained with hoescht (X-axis) and anti-pH3-Ser10 (Y-axis) and analyzed by flow cytometry. (**I**) Western blot analysis of interphase, cycling, and STLC-arrested mitotic cells show similar and background levels of phospho-eIF2 α. Synchronized mitotic cells do not hyper-phosphorylate eIF2α. Sodium arsenite treatment (1 hour, 0.5mM) was as a marker for hyper-phosphorylated eIF2α. Right, quantification of Western blots. (**J**) Principal component analysis of translation initiation site reads from interphase, cycling, and STLC-arrested mitotic cells. Each point is a single biological replicate. Shapes represent different batches. (**K**) A scatter plot shows that the fold change in translation initiation efficiency between cycling (Y-axis) and STLC-arrested (X-axis) relative to interphase cells is correlated. Since the interphase and STLC-arrested samples were prepared in the same batch, we compared interphase to STLC-arrested cells for the manuscript.

**Extended data Fig. 2.**
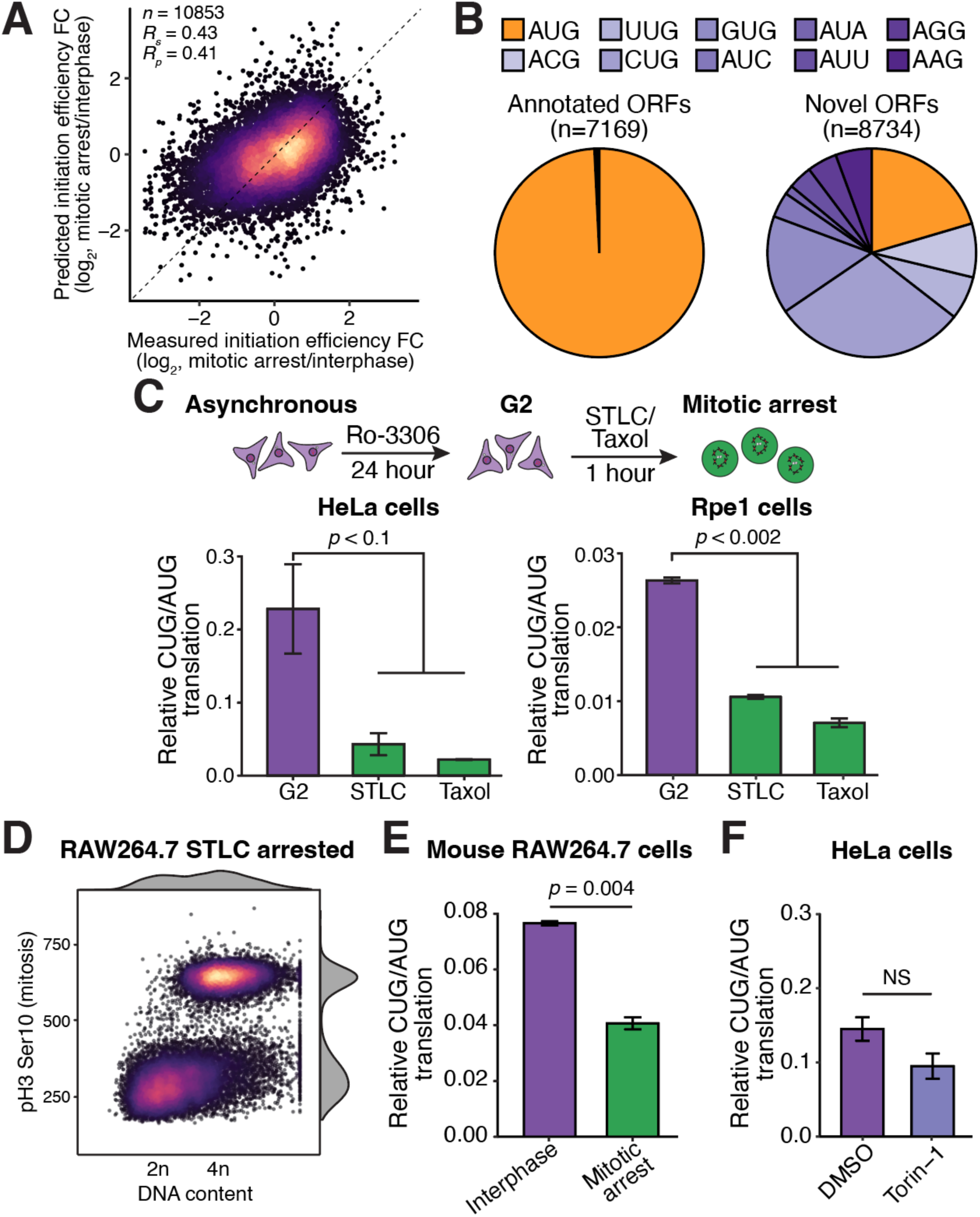
Non-AUG translation is preferentially repressed in mitosis across organisms and cell types. (**A**) Performance of linear regression models to predict translation initiation efficiency fold change between interphase and mitosis. Predictions were concatenated from all folds of held-out data. (**B**) Translation initiation site codon counts from annotated and novel ORFs (extensions, truncations, uORFs, and altORFs). Non-AUG initiation sites only includes near-cognate initiation sites. (**C**) Top, schematic of different cell synchronization strategy. G2 arrested (Ro-3306) cells were released into STLC or taxol media were transfected with mRNA reporters. Bar graph showing that STLC or taxol arrested cells increase translation initiation stringency compared to G2 arrested HeLa (left) and Rpe1 (right) cells. (**D**) Mitotic synchronization efficiency of mouse RAW264.7 cells. X-axis is DNA content based on hoescht intensity and Y-axis is the abundance of the mitotic marker, pH3Ser10. (**E**) Luciferase mRNA reporter assays in synchronized RAW264.7 cells. Error bar represents standard error of the mean, N = 2 biological replicates and Unpaired student’s T-test was used. (**F**) Luciferase mRNA reporter assays HeLa cells treated with Torin-1. Error bar represents standard error of the mean, N = 2 biological replicates and Unpaired student’s T-test was used.

**Extended data Fig. 3.**
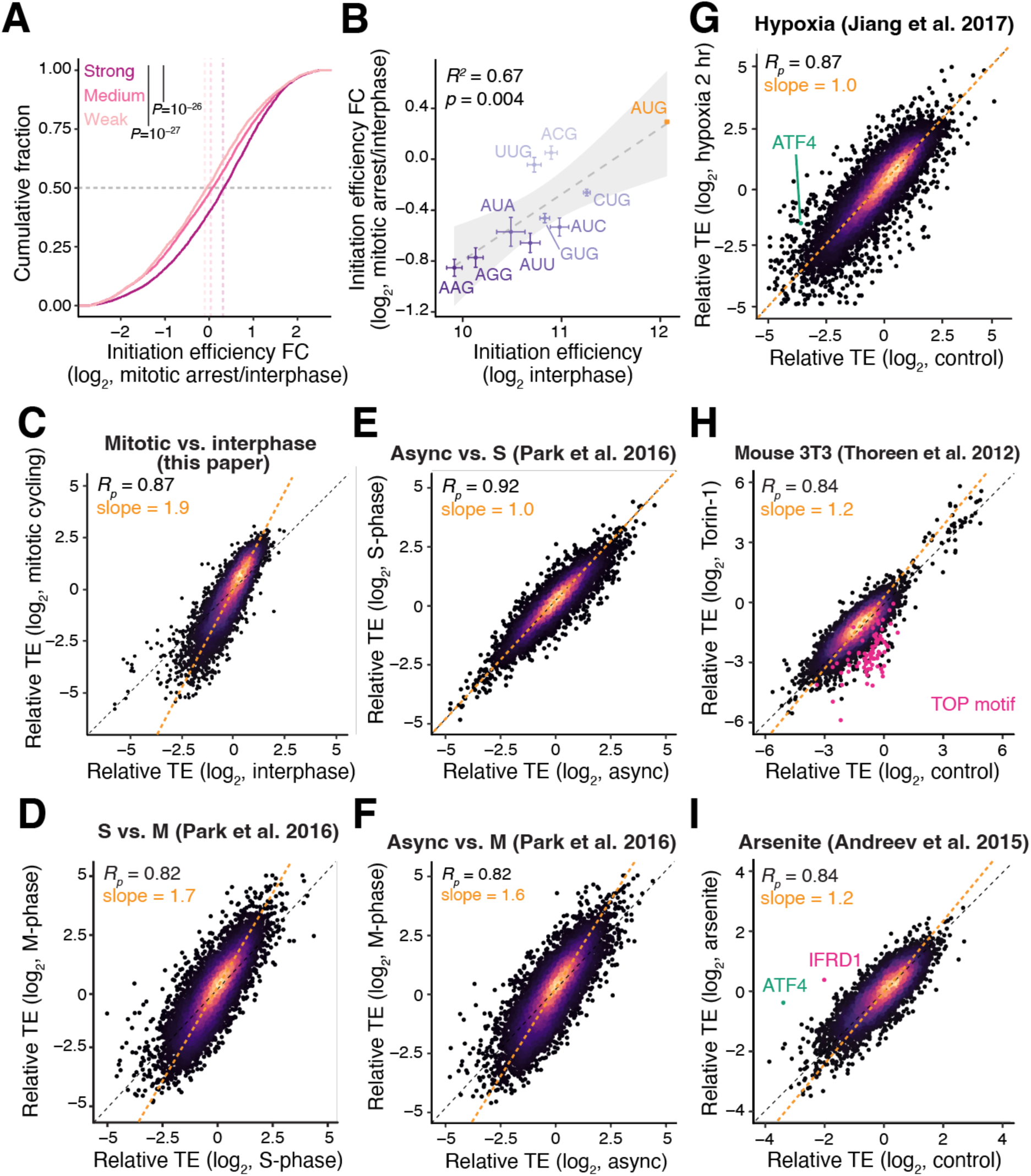
Translation initiation stringency is increased during mitosis. (**A**) Cumulative distribution function (CDF) plot of translation initiation efficiency fold-change between mitosis and interphase for weak, medium, and strong Kozak contexts sites. Strong Kozak contexts are defined as having a -3 G/A and a +4 G, medium contexts contain only either a -3 G/A or +4 G, and weak contexts have neither. (**B**) Scatter plot showing the correlation between interphase translation initiation efficiency (X-axis) and the fold-change between mitotic arrest/interphase (Y-axis). Each point represents the average initiation efficiency or fold change in initiation efficiency between interphase and mitotic arrest. Error bars represent standard error of the mean. The grey shading represents the 95% confidence for the line of best fit. (**C**) Relative translational efficiency correlations between different interphase and cycling mitotic HeLa cells. (**D**) Relative translational efficiency correlations from ^21^, comparing S-phase and mitotically enriched HeLa cells. We note that the synchronization in ^21^ did not involve mitotic shake off so likely contains contaminating interphase/G2 cells in the mitotic prep. (**E**) Relative translational efficiency correlations from ^21^, comparing asynchronous and S-phase cells. and mitotically enriched cells. We note that the synchronization in ^21^ did not involve mitotic shake off so likely contains contaminating interphase/G2 Helacells in the mitotic prep. (**F**) Relative translational efficiency correlations from ^21^, comparing asynchronous and mitotically enriched HeLa cells. We note that the synchronization in ^21^ did not involve mitotic shake off so likely contains contaminating interphase/G2 cells in the mitotic prep. (**G**) Relative translational efficiency correlations between control cells and cells cultured under hypoxic conditions for 2 hours (data from ^32^). (**H**) Relative translational efficiency correlations from ^33^, comparing control and Torin-1 treated mouse 3T3 cells. (**I**) Relative translational efficiency correlations from ^34^, comparing control and arsenite treated HEK293T cells.

**Extended data Fig. 4.**
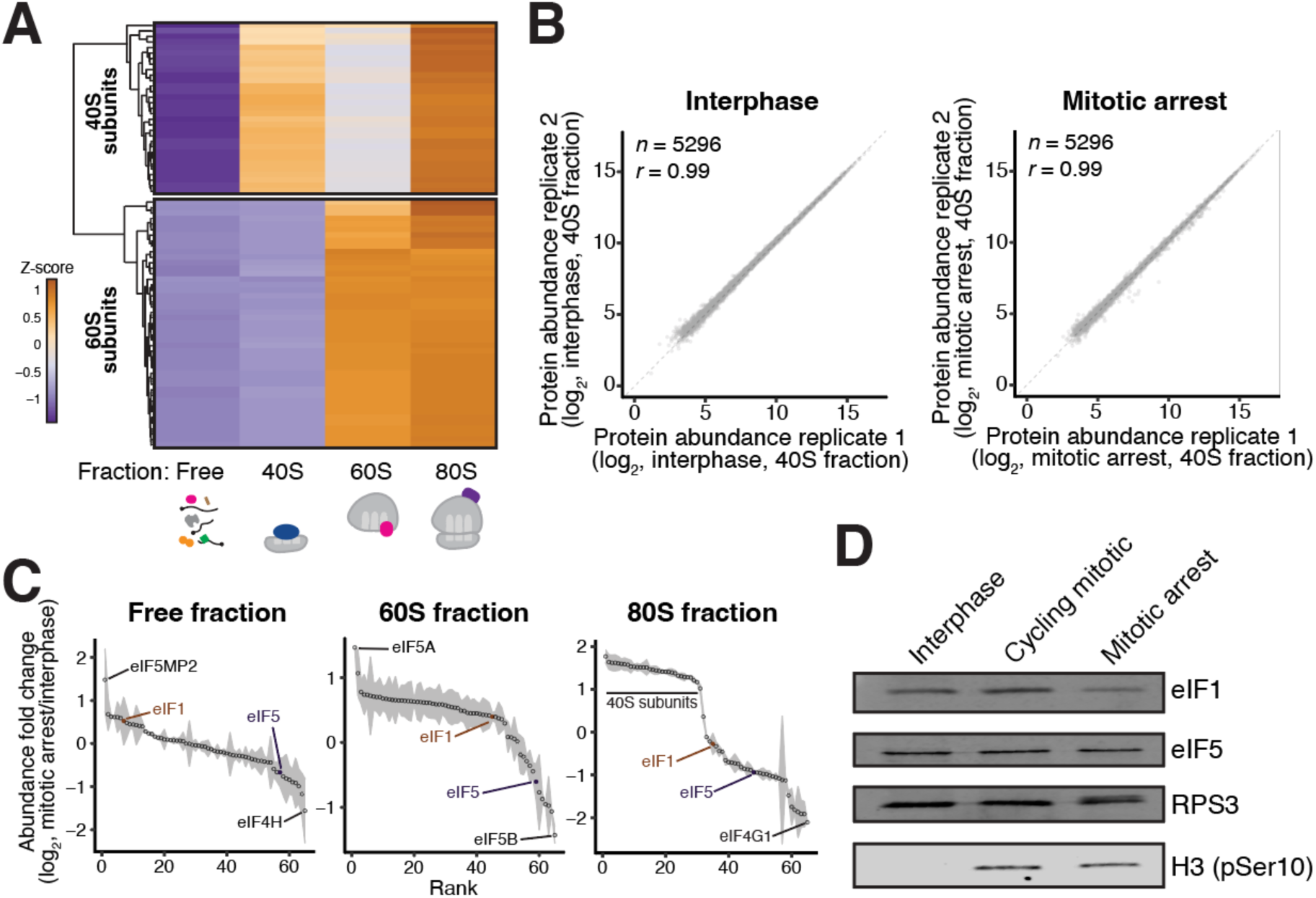
eIF1 preferentially associates with the ribosome during mitosis. (**A**) Heatmap showing successful fractionation of ribosomes and quantitative nature of mass spectrometry from interphase cells. Top cluster represents 40S subunits while bottom represents 60S subunits. Plotted is the average Z-score of 2 biological replicates. (**B**) Biological replicates from interphase (left) and mitotic (right) 40S fraction TMT mass spectrometry experiments. (**C**) Rank-ordered fold-change in the abundance of translation initiation factors and ribosomal proteins in free, 60S, and 80S ribosome fractions. Each point represents the average fold change between 2 biological replicates and the shading is the standard error of the mean. (**D**) Western blot analysis of eIF1 and eIF5 protein expression in interphase, cycling mitotic, and mitotically arrested cells.

**Extended data Fig. 5.**
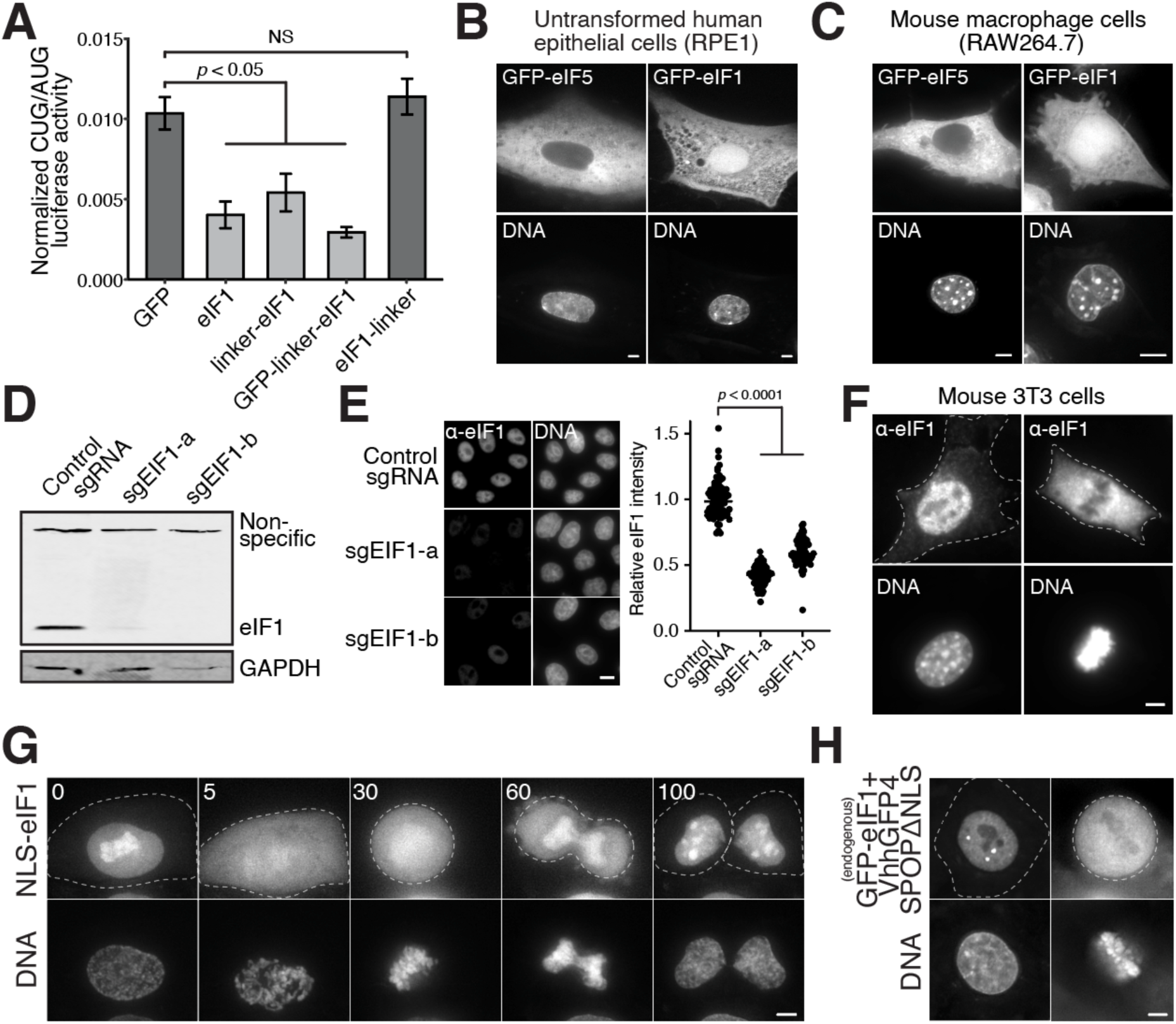
Nuclear eIF1 is released into the cytoplasm during mitosis and meiosis. (**A**) Luciferase reporter assays with co-expressed eIF1 tagged variants. AUG and CUG nano luciferase reporters as described in ^60^, were co-transfected with control AUG firefly luciferase and different eIF1 constructs. N-terminally tagged eIF1 is able to preferentially repress the CUG reporter but C-terminally tagged eIF1 is inactive. N = 3 biological replicates and unpaired student’s T-test was used. (**B**) Localization of GFP-eIF1 and GFP-eIF5 in Rpe1 cells. RPE1 cells were transiently transfected with the GFP-tagged constructs. Scale bar, 5 µm. (**C**) Localization of GFP-eIF1 and GFP-eIF5 in mouse RAW264.7 cells. Scale bar, 5 µm. (**D**) Validation of affinity purified α-eIF1 antibody by Western blotting. Lysates from transient control or eIF1 knockout cells (3-day post infection, using 2 separate sgRNAs targeting eIF1) were analyzed using our affinity purified α-eIF1 antibody. (**E**) Validation of affinity purified α-eIF1 antibody by immunofluorescence. Representative image is shown on the left and quantification from biological replicates shown on the right. Unpaired student’s T-test was used. Scale bar, 10 µm. (**F**) Endogenous eIF1 localization in interphase and mitotic mouse NIH3T3. Dotted line represents cell boundary. Scale bar, 5 µm. (**G**) Live-cell timelapse microscopy of NLS-GFP-eIF1 showing the release of the nuclear fraction of eIF1 into the cytoplasm during mitosis. Numbers indicate time in minutes. Dotted lines represent cell boundaries Scale bar, 5 µm. (**H**) Live-cell imaging of endogenously tagged GFP-eIF1 cells in which the cytoplasmic fraction of eIF1 was eliminated using a cytoplasm specific GFP-degron. The nuclear fraction of eIF1 is released into the cytoplasm during mitosis (right). Dotted lines represent cell boundaries. Scale bar, 5 µm.

**Extended data Fig. 6.**
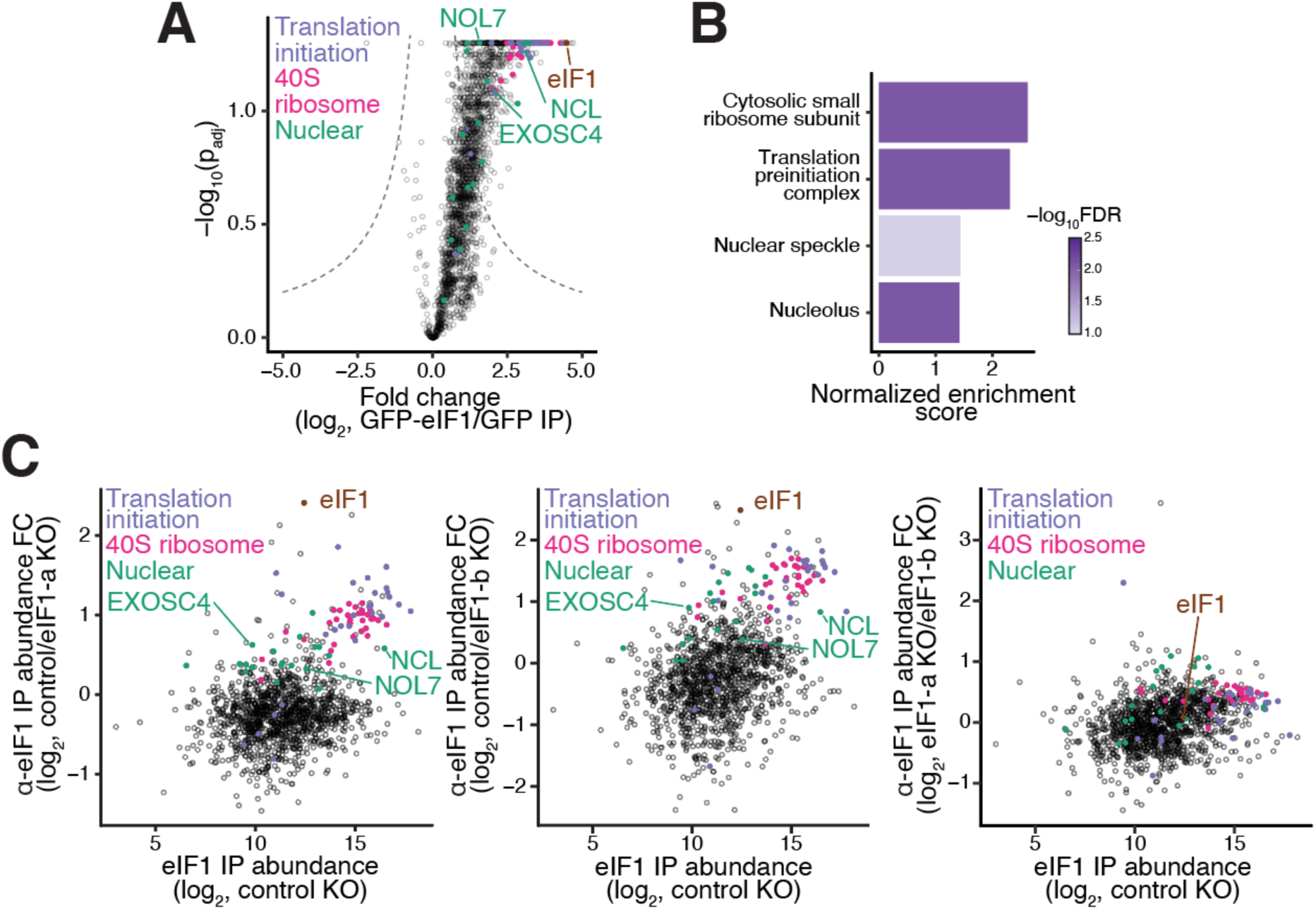
eIF1 interacts with multiple nuclear proteins. (**A**) mEGFP-eIF1 or 3xHA-GFP was immunoprecipitated using anti-GFP and analyzed by quantitative mass spectrometry. Dashed lines indicate the significance threshold. N = 2 biological replicates. (**B**) Gene set enrichment analysis from GFP-eIF1 IP-MS experiment. (**C**) Endogenous eIF1 was immunoprecipitated using α-eIF1 antibody and analyzed by quantitative mass spectrometry. Immunoprecipitations were done from control, and two transient eIF1 knockout interphase cells. The protein abundance from α-eIF1 IP is shown on the X-axis. We note that this abundance is not length normalized. On the Y-axis, we have plotted the abundance fold change between α-eIF1 immunoprecipitations from control and eIF1 knockout cells. Right plot shows lack of change between the two independent eIF1 knockdowns. Selected nuclear factors (see Table S6) are highlighted in green.

**Extended data Fig. 7.**
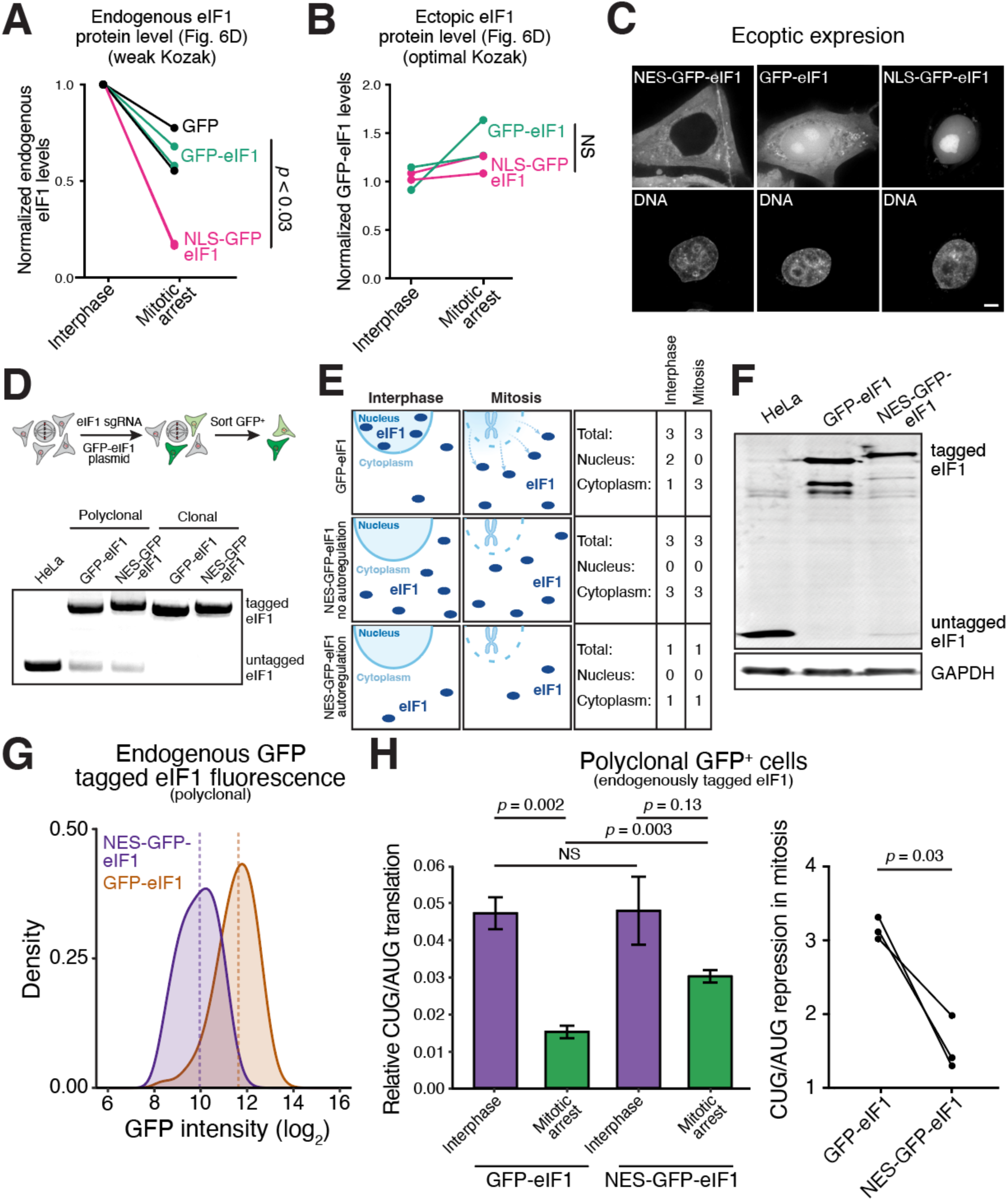
Nuclear/cytoplasmic ratios of initiation factors differentially control translation initiation stringency during mitosis. (A) Quantification of endogenous eIF1 protein levels from Fig. 4D. Overexpressing nuclear eIF1 specifically induces autoregulation in mitosis. (**B**) Quantification of transgenic GFP-eIF1 protein levels from Fig. 4D. Transgenic eIF1 under a strong Kozak context escapes the autoregulation, consistent with translation initiation dependent control. (**C**) Localization of ectopically expressed NES-GFP-eIF1, GFP-eIF1, and NLS-GFP-eIF1. Scale bar, 5 µm. (**D**) Left, schematic outline to generate a polyclonal population of GFP-eIF1 or NES-GFP-eIF1 cells. Right, genotyping PCR of HeLa, polyclonal and monoclonal GFP-eIF1 or NES-GFP-eIF1 cells. (**E**) Right, schematic representation of the role of autoregulation in affecting relative nuclear and cytoplasmic eIF1 levels upon induced nuclear export. Left, hypothetical ratios of eIF1 in the nucleus and cytoplasm with and without autoregulation during interphase and mitosis. Autoregulation during interphase maintains a similar level of cytoplasmic eIF1 upon induced nuclear export. Therefore, this nuclear export selectively depletes nuclear eIF1, while keeping cytoplasmic eIF1 similar. During mitosis, nuclear release of eIF1 doesn’t occur when eIF1 is endogenously tagged with a NES. Therefore, NES-GFP-eIF1 cells have reduced eIF1 activity specifically during mitosis. (**F**) Western blot showing decreased levels of polyclonal NES-GFP-eIF1 protein relative to GFP-eIF1 cells. (**G**) Flow cytometry analysis of the polyclonal population of endogenous GFP-eIF1 and NES-GFP eIF1. (**H**) Luciferase mRNA reporter assays as described in Fig. 4G, except using polyclonal lines.

**Extended data Fig. 8.**
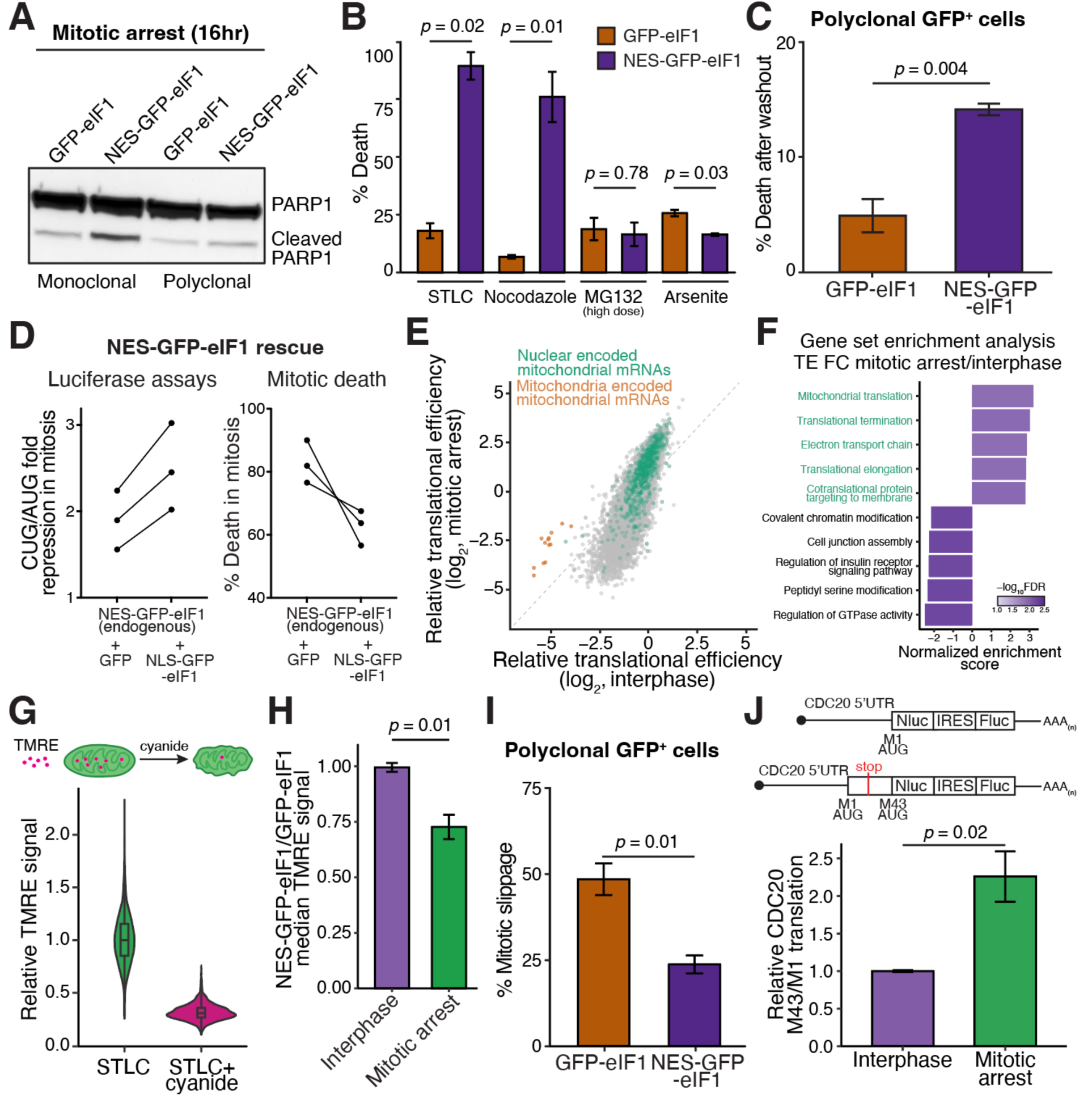
Depleting nuclear eIF1 sensitizes cells to anti-mitotic chemotherapeutics. **(A)** Western blot of PARP1 cleavage of mitotically arrested GFP-eIF1 and NES-GFP-eIF1 cells. Increased PARP1 cleavage in NES-GFP-eIF1 cells is indicative of apoptosis. (**B**) Death in the presence of various drugs measured by propidium iodide staining. Note that the high concentration of MG132 used in this experiment interphase cells do not enter mitosis therefore does not enrich for mitotic cells. Error bar represents standard error of the mean, N = 2 biological replicates and Unpaired student’s T-test was used. (**C**) STLC washout assays as described in Fig. 5C, except using polyclonal lines. (**D**) Rescue of translation initiation stringency and mitotic death in endogenous NES-GFP-eIF1 cells through ectopic expression of NLS-GFP-eIF1. Left, luciferase mRNA reporter assays, plotting the repression of CUG/AUG ratio in mitosis in cells lacking nuclear eIF1 (endogenous NES-GFP-eIF1 + GFP, first column) and with nuclear eIF1 (endogenous NES-GFP-eIF1 + NLS-GFP-eIF1, second column). Right, mitotic death in taxol using live-cell imaging in cells with and without nuclear eIF1. (**E**) Mitochondrial mRNAs, encoded from both the mitochondria and nucleus, are preferentially translated during mitosis. Removal of nuclear eIF1 is predicted to shift the slope of the plot closer to 1, therefore reducing the translation of mitochondrial mRNAs during mitosis. (**F**) Gene set enrichment analysis using change in translational efficiency between mitotically arrested and interphase cells. (**G**) Positive control experiment to show that TMRE is staining active mitochondria. Addition of cyanide poisons the electron transport chain, resulting in mitochondrial depolarization. TMRE signal is quantified using Cell Profiler. (**H**) Median TMRE signal in 3 biological replicates. Error bar represents standard error of the mean and unpaired student’s T-test was used. (**I**) Mitotic slippage assays as described in Fig. 5F, except using polyclonal lines. (**J**) Luciferase mRNA reporter assays to assess the relative translation of CDC20 M1 and M43 translation initiation sites in interphase and mitotically arrested cells. CDC20 M43 reporter is preferentially translated relative to CDC20 M1 reporter in mitosis, consistent with nuclear eIF1 release promoting leaky scanning in mitosis. Error bar represents standard error of the mean, N = 3 biological replicates and unpaired student’s T-test used for this data.

## References and Notes

1 Wright, B. W., Yi, Z., Weissman, J. S. & Chen, J. The dark proteome: translation from noncanonical open reading frames. Trends Cell Biol 32, 243–258 (2022). 10.1016/j.tcb.2021.10.010

2 Conti, M. & Kunitomi, C. A genome-wide perspective of the maternal mRNA translation program during oocyte development. Semin Cell Dev Biol (2023). 10.1016/j.semcdb.2023.03.003

3 Vastenhouw, N. L., Cao, W. X. & Lipshitz, H. D. The maternal-to-zygotic transition revisited. Development 146 (2019). 10.1242/dev.161471

4 Palozola, K. C., et al. Mitotic transcription and waves of gene reactivation during mitotic exit. Science 358, 119–122 (2017). 10.1126/science.aal4671

5 Shin, C. & Manley, J. L. The SR protein SRp38 represses splicing in M phase cells. Cell 111, 407–417 (2002). 10.1016/s0092-8674(02)01038-3

6 Parsons, G. G. & Spencer, C. A. Mitotic repression of RNA polymerase II transcription is accompanied by release of transcription elongation complexes. Mol Cell Biol 17, 5791–5802 (1997). 10.1128/MCB.17.10.5791

7 McAinsh, A. D. & Kops, G. Principles and dynamics of spindle assembly checkpoint signalling. Nat Rev Mol Cell Biol (2023). 10.1038/s41580-023-00593-z

8 Lara-Gonzalez, P., Pines, J. & Desai, A. Spindle assembly checkpoint activation and silencing at kinetochores. Semin Cell Dev Biol 117, 86–98 (2021). 10.1016/j.semcdb.2021.06.009

9 Musacchio, A. The Molecular Biology of Spindle Assembly Checkpoint Signaling Dynamics. Curr Biol 25, R1002–1018 (2015). 10.1016/j.cub.2015.08.051

10 Musacchio, A. & Salmon, E. D. The spindle-assembly checkpoint in space and time. Nat Rev Mol Cell Biol 8, 379–393 (2007). 10.1038/nrm2163

11 Skoufias, D. A., et al. S-trityl-L-cysteine is a reversible, tight binding inhibitor of the human kinesin Eg5 that specifically blocks mitotic progression. J Biol Chem 281, 17559–17569 (2006). 10.1074/jbc.M511735200

12 Lindqvist, L. M., et al. Translation inhibitors induce cell death by multiple mechanisms and Mcl-1 reduction is only a minor contributor. Cell Death Dis 3, e409 (2012). 10.1038/cddis.2012.149

13 Mouilleron, H., Delcourt, V. & Roucou, X. Death of a dogma: eukaryotic mRNAs can code for more than one protein. Nucleic Acids Res 44, 14–23 (2016). 10.1093/nar/gkv1218

14 Andreev, D. E., et al. Non-AUG translation initiation in mammals. Genome Biol 23, 111 (2022). 10.1186/s13059-022-02674-2

15 Zhang, P., et al. Genome-wide identification and differential analysis of translational initiation. Nat Commun 8, 1749 (2017). 10.1038/s41467-017-01981-8

16 Coldwell, M. J., et al. Phosphorylation of eIF4GII and 4E-BP1 in response to nocodazole treatment: a reappraisal of translation initiation during mitosis. Cell Cycle 12, 3615–3628 (2013). 10.4161/cc.26588

17 Yang, X., et al. Human BAG-1/RAP46 protein is generated as four isoforms by alternative translation initiation and overexpressed in cancer cells. Oncogene 17, 981–989 (1998). 10.1038/sj.onc.1202032

18 Loughran, G., et al. Unusually efficient CUG initiation of an overlapping reading frame in POLG mRNA yields novel protein POLGARF. Proc Natl Acad Sci U S A 117, 24936–24946 (2020). 10.1073/pnas.2001433117

19 Hann, S. R., King, M. W., Bentley, D. L., Anderson, C. W. & Eisenman, R. N. A non-AUG translational initiation in c-myc exon 1 generates an N-terminally distinct protein whose synthesis is disrupted in Burkitt’s lymphomas. Cell 52, 185–195 (1988). 10.1016/0092-8674(88)90507-7

20 Tanenbaum, M. E., Stern-Ginossar, N., Weissman, J. S. & Vale, R. D. Regulation of mRNA translation during mitosis. Elife 4 (2015). 10.7554/eLife.07957

21 Park, J. E., Yi, H., Kim, Y., Chang, H. & Kim, V. N. Regulation of Poly(A) Tail and Translation during the Somatic Cell Cycle. Mol Cell 62, 462–471 (2016). 10.1016/j.molcel.2016.04.007

22 Saris, C. J., Domen, J. & Berns, A. The pim-1 oncogene encodes two related protein-serine/threonine kinases by alternative initiation at AUG and CUG. EMBO J 10, 655–664 (1991). 10.1002/j.1460-2075.1991.tb07994.x

23 Zhang, X., et al. Translational control of the cytosolic stress response by mitochondrial ribosomal protein L18. Nat Struct Mol Biol 22, 404–410 (2015). 10.1038/nsmb.3010

24 Jin, X., Turcott, E., Englehardt, S., Mize, G. J. & Morris, D. R. The two upstream open reading frames of oncogene mdm2 have different translational regulatory properties. J Biol Chem 278, 25716–25721 (2003). 10.1074/jbc.M300316200

25 Luke J. Fulcher, T. S., Ian Gibbs-Seymour, Francis A. Barr. MDM2 acts as a timer reporting the length of mitosis. biorxiv (2023). 10.1101/2023.05.26.542398

26 Li, J., et al. Proteome-wide mapping of short-lived proteins in human cells. Mol Cell 81, 4722–4735 e4725 (2021). 10.1016/j.molcel.2021.09.015

27 Timms, R. T., et al. A glycine-specific N-degron pathway mediates the quality control of protein N-myristoylation. Science 365 (2019). 10.1126/science.aaw4912

28 Kozak, M. Point mutations define a sequence flanking the AUG initiator codon that modulates translation by eukaryotic ribosomes. Cell 44, 283–292 (1986). 10.1016/0092-8674(86)90762-2

29 Kozak, M. An analysis of 5’-noncoding sequences from 699 vertebrate messenger RNAs. Nucleic Acids Res 15, 8125–8148 (1987). 10.1093/nar/15.20.8125

30 Ingolia, N. T., Lareau, L. F. & Weissman, J. S. Ribosome profiling of mouse embryonic stem cells reveals the complexity and dynamics of mammalian proteomes. Cell 147, 789–802 (2011). 10.1016/j.cell.2011.10.002

31 Ingolia, N. T. Ribosome Footprint Profiling of Translation throughout the Genome. Cell 165, 22–33 (2016). 10.1016/j.cell.2016.02.066

32 Jiang, Z., et al. Ribosome profiling reveals translational regulation of mammalian cells in response to hypoxic stress. BMC Genomics 18, 638 (2017). 10.1186/s12864-017-3996-8

33 Thoreen, C. C., et al. A unifying model for mTORC1-mediated regulation of mRNA translation. Nature 485, 109–113 (2012). 10.1038/nature11083

34 Andreev, D. E., et al. Translation of 5’ leaders is pervasive in genes resistant to eIF2 repression. Elife 4, e03971 (2015). 10.7554/eLife.03971

35 Hinnebusch, A. G. & Lorsch, J. R. The mechanism of eukaryotic translation initiation: new insights and challenges. Cold Spring Harb Perspect Biol 4 (2012). 10.1101/cshperspect.a011544

36 Pestova, T. V. & Kolupaeva, V. G. The roles of individual eukaryotic translation initiation factors in ribosomal scanning and initiation codon selection. Genes Dev 16, 2906–2922 (2002). 10.1101/gad.1020902

37 Ivanov, I. P., Loughran, G., Sachs, M. S. & Atkins, J. F. Initiation context modulates autoregulation of eukaryotic translation initiation factor 1 (eIF1). Proc Natl Acad Sci U S A 107, 18056–18060 (2010). 10.1073/pnas.1009269107

38 Cheung, Y. N., et al. Dissociation of eIF1 from the 40S ribosomal subunit is a key step in start codon selection in vivo. Genes Dev 21, 1217–1230 (2007). 10.1101/gad.1528307

39 Manjunath, H., et al. Suppression of Ribosomal Pausing by eIF5A Is Necessary to Maintain the Fidelity of Start Codon Selection. Cell Rep 29, 3134–3146 e3136 (2019). 10.1016/j.celrep.2019.10.129

40 Loughran, G., Sachs, M. S., Atkins, J. F. & Ivanov, I. P. Stringency of start codon selection modulates autoregulation of translation initiation factor eIF5. Nucleic Acids Res 40, 2898–2906 (2012). 10.1093/nar/gkr1192

41 Ivanov, I. P., et al. Evolutionarily conserved inhibitory uORFs sensitize Hox mRNA translation to start codon selection stringency. Proc Natl Acad Sci U S A 119 (2022). 10.1073/pnas.2117226119

42 Llacer, J. L., et al. Translational initiation factor eIF5 replaces eIF1 on the 40S ribosomal subunit to promote start-codon recognition. Elife 7 (2018). 10.7554/eLife.39273

43 Tang, L., et al. Competition between translation initiation factor eIF5 and its mimic protein 5MP determines non-AUG initiation rate genome-wide. Nucleic Acids Res 45, 11941–11953 (2017). 10.1093/nar/gkx808

44 Pisareva, V. P. & Pisarev, A. V. eIF5 and eIF5B together stimulate 48S initiation complex formation during ribosomal scanning. Nucleic Acids Res 42, 12052–12069 (2014). 10.1093/nar/gku877

45 Ghosh, A., Jindal, S., Bentley, A. A., Hinnebusch, A. G. & Komar, A. A. Rps5-Rps16 communication is essential for efficient translation initiation in yeast S. cerevisiae. Nucleic Acids Res 42, 8537–8555 (2014). 10.1093/nar/gku550

46 Nanda, J. S., et al. eIF1 controls multiple steps in start codon recognition during eukaryotic translation initiation. J Mol Biol 394, 268–285 (2009). 10.1016/j.jmb.2009.09.017

47 Hornbeck, P. V. et al. PhosphoSitePlus, 2014: mutations, PTMs and recalibrations. Nucleic Acids Res 43, D512–520 (2015). 10.1093/nar/gku1267

48 Petrone, A., Adamo, M. E., Cheng, C. & Kettenbach, A. N. Identification of Candidate Cyclin-dependent kinase 1 (Cdk1) Substrates in Mitosis by Quantitative Phosphoproteomics. Mol Cell Proteomics 15, 2448–2461 (2016). 10.1074/mcp.M116.059394

49 Bohnsack, M. T., et al. Exp5 exports eEF1A via tRNA from nuclei and synergizes with other transport pathways to confine translation to the cytoplasm. EMBO J 21, 6205–6215 (2002). 10.1093/emboj/cdf613

50 Cho, N. H., et al. OpenCell: Endogenous tagging for the cartography of human cellular organization. Science 375, eabi6983 (2022). 10.1126/science.abi6983

51 Fijalkowska, D., et al. eIF1 modulates the recognition of suboptimal translation initiation sites and steers gene expression via uORFs. Nucleic Acids Res 45, 7997–8013 (2017). 10.1093/nar/gkx469

52 Zhou, F., Zhang, H., Kulkarni, S. D., Lorsch, J. R. & Hinnebusch, A. G. eIF1 discriminates against suboptimal initiation sites to prevent excessive uORF translation genome-wide. RNA 26, 419–438 (2020). 10.1261/rna.073536.119

53 Weaver, B. A. How Taxol/paclitaxel kills cancer cells. Mol Biol Cell 25, 2677–2681 (2014). 10.1091/mbc.E14-04-0916

54 Crowley, L. C., et al. Measuring Cell Death by Propidium Iodide Uptake and Flow Cytometry. Cold Spring Harb Protoc 2016 (2016). 10.1101/pdb.prot087163

55 Chaitanya, G. V., Steven, A. J. & Babu, P. P. PARP-1 cleavage fragments: signatures of cell-death proteases in neurodegeneration. Cell Commun Signal 8, 31 (2010). 10.1186/1478-811X-8-31

56 Bock, F. J. & Tait, S. W. G. Mitochondria as multifaceted regulators of cell death. Nat Rev Mol Cell Biol 21, 85–100 (2020). 10.1038/s41580-019-0173-8

57 Ghelli Luserna di Rora, A., Martinelli, G. & Simonetti, G. The balance between mitotic death and mitotic slippage in acute leukemia: a new therapeutic window? J Hematol Oncol 12, 123 (2019). 10.1186/s13045-019-0808-4

58 Gascoigne, K. E. & Taylor, S. S. Cancer cells display profound intra- and interline variation following prolonged exposure to antimitotic drugs. Cancer Cell 14, 111–122 (2008). 10.1016/j.ccr.2008.07.002

59 Tsang, M. J. & Cheeseman, I. M. Alternative CDC20 translational isoforms tune mitotic arrest duration. Nature 617, 154–161 (2023). 10.1038/s41586-023-05943-7

60 Kearse, M. G., et al. Ribosome queuing enables non-AUG translation to be resistant to multiple protein synthesis inhibitors. Genes Dev 33, 871–885 (2019). 10.1101/gad.324715.119

61 Singh, C. R., et al. Human oncoprotein 5MP suppresses general and repeat-associated non-AUG translation via eIF3 by a common mechanism. Cell Rep 36, 109376 (2021). 10.1016/j.celrep.2021.109376

62 Clemm von Hohenberg, K., et al. Cyclin B/CDK1 and Cyclin A/CDK2 phosphorylate DENR to promote mitotic protein translation and faithful cell division. Nat Commun 13, 668 (2022). 10.1038/s41467-022-28265-0

63 Eisenberg, A. R., et al. Translation Initiation Site Profiling Reveals Widespread Synthesis of Non-AUG-Initiated Protein Isoforms in Yeast. Cell Syst 11, 145–160 e145 (2020). 10.1016/j.cels.2020.06.011

64 Guenther, U. P., et al. The helicase Ded1p controls use of near-cognate translation initiation codons in 5’ UTRs. Nature 559, 130–134 (2018). 10.1038/s41586-018-0258-0

65 Stumpf, C. R., Moreno, M. V., Olshen, A. B., Taylor, B. S. & Ruggero, D. The translational landscape of the mammalian cell cycle. Mol Cell 52, 574–582 (2013). 10.1016/j.molcel.2013.09.018

66 Fan, H. & Penman, S. Regulation of protein synthesis in mammalian cells. II. Inhibition of protein synthesis at the level of initiation during mitosis. J Mol Biol 50, 655–670 (1970). 10.1016/0022-2836(70)90091-4

67 Zecha, J., et al. Peptide Level Turnover Measurements Enable the Study of Proteoform Dynamics. Mol Cell Proteomics 17, 974–992 (2018). 10.1074/mcp.RA118.000583

68 Eden, E., et al. Proteome half-life dynamics in living human cells. Science 331, 764–768 (2011). 10.1126/science.1199784

69 Fujiwara, T., et al. Cytokinesis failure generating tetraploids promotes tumorigenesis in p53-null cells. Nature 437, 1043–1047 (2005). 10.1038/nature04217

70 Huang, H. C., Shi, J., Orth, J. D. & Mitchison, T. J. Evidence that mitotic exit is a better cancer therapeutic target than spindle assembly. Cancer Cell 16, 347–358 (2009). 10.1016/j.ccr.2009.08.020

71 Martin, M. Cutadapt removes adapter sequences from high-throughput sequencing reads. EMBnet.journal 17.1 (2011).

72 Dobin, A., et al. STAR: ultrafast universal RNA-seq aligner. Bioinformatics 29, 15–21 (2013). 10.1093/bioinformatics/bts635

73 Wu, X. & Bartel, D. P. kpLogo: positional k-mer analysis reveals hidden specificity in biological sequences. Nucleic Acids Res 45, W534–W538 (2017). 10.1093/nar/gkx323

74 Anders, S., Pyl, P. T. & Huber, W. HTSeq--a Python framework to work with high-throughput sequencing data. Bioinformatics 31, 166-169 (2015). 10.1093/bioinformatics/btu638

75 Love, M. I., Huber, W. & Anders, S. Moderated estimation of fold change and dispersion for RNA-seq data with DESeq2. Genome Biol 15, 550 (2014). 10.1186/s13059-014-0550-8

76 Rath, S., et al. MitoCarta3.0: an updated mitochondrial proteome now with sub-organelle localization and pathway annotations. Nucleic Acids Res 49, D1541–D1547 (2021). 10.1093/nar/gkaa1011

77 Thul, P. J., et al. A subcellular map of the human proteome. Science 356 (2017). 10.1126/science.aal3321

78 Zhang, S., Hu, H., Jiang, T., Zhang, L. & Zeng, J. TITER: predicting translation initiation sites by deep learning. Bioinformatics 33, i234–i242 (2017). 10.1093/bioinformatics/btx247

79 Gleason, A. C., Ghadge, G., Sonobe, Y. & Roos, R. P. Kozak Similarity Score Algorithm Identifies Alternative Translation Initiation Codons Implicated in Cancers. Int J Mol Sci 23 (2022). 10.3390/ijms231810564

80 Subramanian, A., et al. Gene set enrichment analysis: a knowledge-based approach for interpreting genome-wide expression profiles. Proc Natl Acad Sci U S A 102, 15545–15550 (2005). 10.1073/pnas.0506580102

81 Cheeseman, I. M. & Desai, A. A combined approach for the localization and tandem affinity purification of protein complexes from metazoans. Sci STKE 2005, pl1 (2005). 10.1126/stke.2662005pl1

82 Kall, L., Canterbury, J. D., Weston, J., Noble, W. S. & MacCoss, M. J. Semi-supervised learning for peptide identification from shotgun proteomics datasets. Nat Methods 4, 923–925 (2007). 10.1038/nmeth1113

83 Dang, C. V. & Lee, W. M. Identification of the human c-myc protein nuclear translocation signal. Mol Cell Biol 8, 4048–4054 (1988). 10.1128/mcb.8.10.4048-4054.1988

84 Kosugi, S., et al. Six classes of nuclear localization signals specific to different binding grooves of importin alpha. J Biol Chem 284, 478–485 (2009). 10.1074/jbc.M807017200

85 Gomez Corredor, A. & Archambault, D. The bovine immunodeficiency virus Rev protein: identification of a novel nuclear import pathway and nuclear export signal among retroviral Rev/Rev-like proteins. J Virol 86, 4892–4905 (2012). 10.1128/JVI.05132-11

86 Shin, Y. J., et al. Nanobody-targeted E3-ubiquitin ligase complex degrades nuclear proteins. Sci Rep 5, 14269 (2015). 10.1038/srep14269

